# Integrative Network Analysis Reveals Novel Moderators of Aβ-Tau Interaction in Alzheimer’s Disease

**DOI:** 10.1101/2024.06.14.599092

**Authors:** Akihiro Kitani, Yusuke Matsui

## Abstract

**Background:** Although interactions between amyloid-beta and tau proteins have been implicated in Alzheimer’s disease (AD), the precise mechanisms by which these interactions contribute to disease progression are not yet fully understood. Moreover, despite the growing application of deep learning in various biomedical fields, its application in integrating networks to analyze disease mechanisms in AD research remains limited. In this study, we employed BIONIC, a deep learning-based network integration method, to integrate proteomics and protein–protein interaction data, with an aim to uncover factors that moderate the effects of the Aβ-tau interaction on mild cognitive impairment (MCI) and early-stage AD.

**Methods:** Proteomic data from the ROSMAP cohort were integrated with protein–protein interaction (PPI) data using a Deep Learning-based model. Linear regression analysis was applied to histopathological and gene expression data, and mutual information was used to detect moderating factors. Statistical significance was determined using the Benjamini-Hochberg correction (p < 0.05).

**Results:** Our results suggested that astrocytes and GPNMB+ microglia moderate the Aβ-tau interaction. Based on linear regression with histopathological and gene expression data, GFAP and IBA1 levels and *GPNMB* gene expression positively contributed to the interaction of tau with Aβ in non-dementia cases, replicating the results of the network analysis.

**Conclusions:** These findings indicate that GPNMB+ microglia moderate the Aβ-tau interaction in early AD and therefore are a novel therapeutic target. To facilitate further research, we have made the integrated network available as a visualization tool for the scientific community (URL: https://igcore.cloud/GerOmics/AlzPPMap).

## Background

Alzheimer’s disease (AD) is the most common form of dementia, characterized by extracellular amyloid-beta (Aβ) plaques and intracellular tau protein tangles, which are believed to be central to its pathology.[1–3] Recent studies have reported an interaction between Aβ and tau proteins in the pathogenesis of AD, contributing to disease progression.[4–9] For instance, Lee et al.[7] revealed that Aβ-tau interactions are associated with the propagation of tau. In practice, lecanemab, an anti-Aβ antibody, which has been shown to slow cognitive decline in patients with early-stage AD, reduces total tau protein and P-Tau181 levels in cerebrospinal fluid.[10] This suggests that Aβ contributes to the effect of tau in AD. Moreover, several phenotypes are associated with the Aβ-tau interaction, such as astrocyte reactivity, blood pressure, vascular burden, and microglia[11–15]; however, the detailed mechanisms are not fully understood. Various cellular perturbations have been identified through single-nucleus RNA sequencing (snRNA-seq) in AD.[16–22] For example, a recent snRNA-seq study[17] revealed the presence of Trem2-dependent disease-associated microglia (DAM) and a unique Serpina3n+C4b+ reactive oligodendrocyte population in 5XFAD mice. In human AD, distinct glial phenotypes including *IRF8*-driven reactive microglia and oligodendrocytes with impaired myelination and metabolic adaptation to neuronal degeneration have been observed. However, the specific cell types and effects of Aβ-tau interactions are yet to be elucidated.

Network analyses based on gene or protein co-expression have revealed gene groups and associated pathways correlated with pathological and clinical factors in AD.[23–25] Additionally, network integration methods based on multi-omics and other data types have been developed.[26–29] By integrating both physical and functional interactions simultaneously, network integration analyses can identify detailed molecular relationships. However, this approach has largely been applied to cancer. AD is a complex disease involving various pathologies, and network integration is a promising approach for determining comprehensive physical and functional interactions involving Aβ and tau in AD.

In this study, we investigated moderating factors in the Aβ-tau interaction using a systematic network integration approach. The aim of this study was to elucidate the cell types and moderating factors in the interaction between Aβ and tau proteins in early-stage AD. Figure 1 shows the graphical abstract of the study. First, we integrated the proteomics and protein–protein interaction (PPI) networks by using BIONIC,[30] a network integration algorithm. In the downstream analysis, glial cell-related proteins, including those associated with microglia and astrocytes, were enriched around *APP* and *MAPT* subnetworks, which encode the precursors of Aβ and tau, respectively. Further analyses of microglial subtypes revealed enrichment in GPNMB+ microglia. As a validation, linear regression analysis based on histopathological and gene expression data showed that interactions between Aβ and IBA1, GFAP, and *GPNMB*, were positively related to tau levels. Our findings highlight that deep learning-based network integration can serve as a novel framework for elucidating complex disease mechanisms. The integrated network, made available as a visualization tool for the scientific community, not only enhances our understanding of the Aβ-tau interaction but also provides a basis for generating experimental validation ideas and developing more effective treatments for AD. By leveraging this tool, researchers can gain insights into the intricate molecular interactions at play in AD, fostering the discovery of new therapeutic targets.

**Fig. 1.**
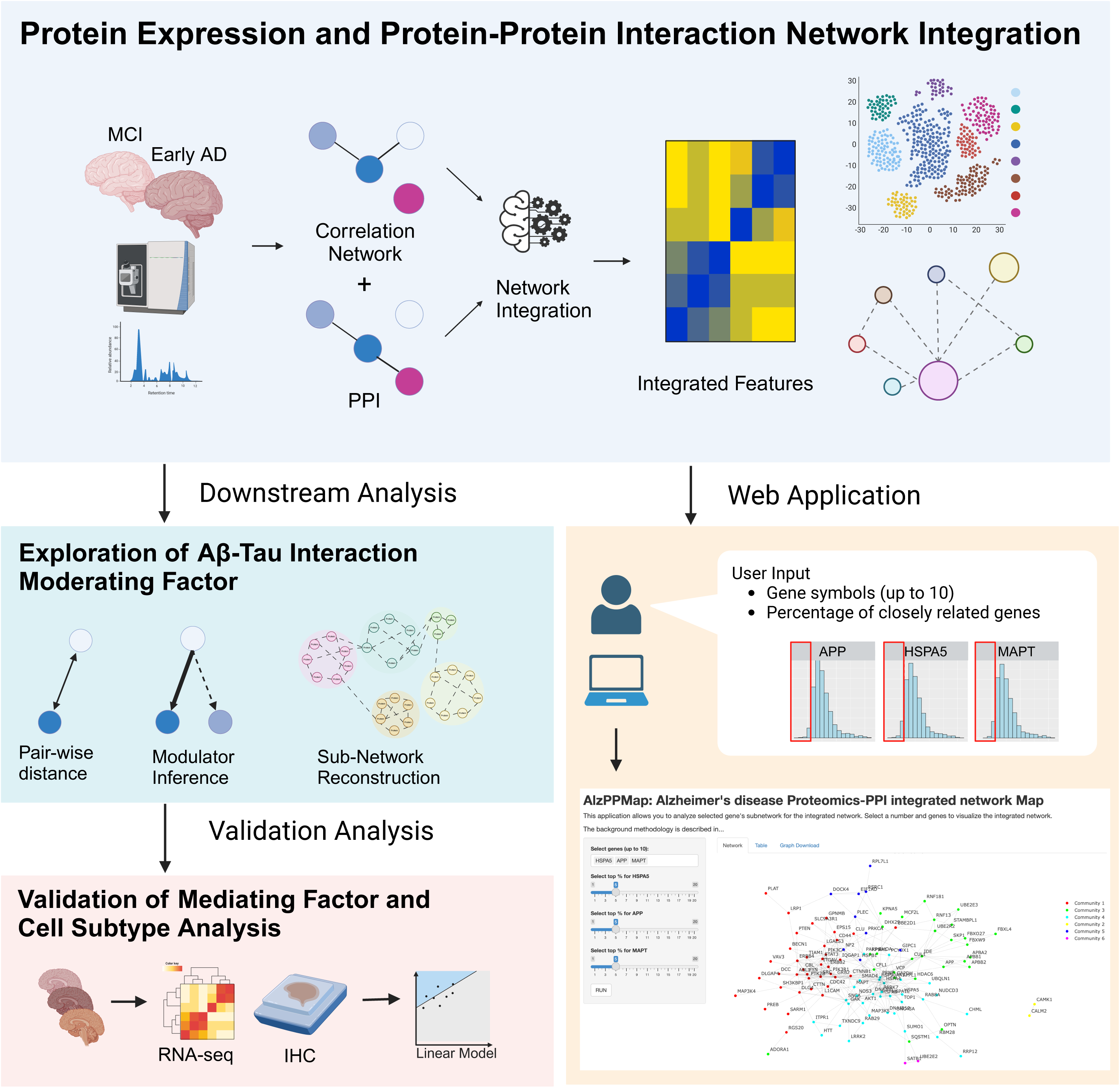
A graphical abstract of the study This study consists of the following four steps: 1. Integration of proteomics and protein–protein interaction (PPI) data, 2. Downstream analysis: Identification of moderators involving APP and MAPT, and construction of subnetworks, 3. Validation using pathological histological evaluation data, 4. Development of a user-friendly web application. In addition to step 2 conducted in this study, users can also construct and visualize subnetworks of genes of interest (including interactions between multiple genes).

## Methods

### Proteomics data from the ROSMAP cohort

For the proteomic analysis, ROSMAP tandem mass tag (TMT) proteomics data were obtained. TMT proteomics is a technique for the simultaneous quantitative analysis of protein expression in multiple samples using LC-MS/MS and is commonly employed in AD research.[24,25] The ROSMAP data set included TMT proteomics data for 400 cases with differences in cognitive status (Alzheimer’s dementia, other dementia, MCI, or no impairment). Data normalization and batch correction were conducted according to previous described procedures.[31] Four outlier samples were excluded from subsequent analyses based on a principal component analysis. In this study, ROSMAP TMT data for MCI and early AD cases were included. Early AD was defined according to previously reported criteria: meeting the NIA-AA core clinical criteria for probable AD dementia and having a global CDR score of 0.5 to 1.0 and a CDR memory box score of 0.5 or greater at screening and baseline.[10] Additionally, the MMSE effectively discriminates between CDR stages 0.5, 1, 2, and 3 but performs poorly in the separation of CDR stages zero and 0.5. The previously established MMSE ranges were 30 for no, 26–29 for questionable, 21–25 for mild, 11–20 for moderate, and 0–10 for severe dementia.[32] We defined early AD as having an MMSE score of ≥21.

### Proteomics and PPI data integration using a Graph Attention Network-based method

To integrate the proteomics and PPI data, total values were calculated for each gene symbol in the proteomics dataset. Pearson correlation coefficients were derived from the proteomics data, retaining only data with Benjamini-Hochberg (BH)-corrected p-values of <0.05. Normalization was performed to set maximum values to 1 and minimum values to 0. For PPI, STRING[33] filtering was used to obtain physical interactions within *Homo sapiens* with confidence scores exceeding 0.7, and normalization was performed to set the maximum values to 1 and minimum values to 0. Proteomic and PPI data were integrated using a recently reported method, BIONIC,[30] which integrates multiple biological networks while preserving information using graph attention networks.[34] Default values were used for the BIONIC hyperparameters. For both proteomics and PPI data, 4,787 proteins common to both datasets were retained. After integrating the two networks, 512-dimensional features were output for each protein.

### Clustering analysis and dimension reduction using features derived from BIONIC

Considering that the data derived from neural networks are nonlinear, clustering was performed following a method similar to that used in snRNA-Seq analyses.[35] After feature calculation using BIONIC and subsequent scaling to Z-scores, Euclidean distances were computed to create a distance matrix. Then, setting k = 25, the k-nearest neighbor method was used to calculate an adjacency matrix and form a graph structure. This graph structure was then subjected to clustering using the Louvain method[36] (resolution = 1) to classify each protein into clusters.

### Cluster centroid analysis using BIONIC-derived features

After implementing BIONIC to evaluate the networks derived from proteomic and PPI data, a clustering analysis of proteins was performed. To analyze the relationships between clusters, the average of the feature values for each cluster was calculated as its centroid. Pearson correlation coefficients and Euclidean distances between the clusters were calculated. Furthermore, the network of clusters was visualized using the reciprocal values of the Euclidean distances as the weights of the edges.

### Cell type enrichment analysis based on human brain and microglia snRNA-seq data

For a cell type enrichment analysis, publicly available human brain[37] and microglia[21,22] snRNA-seq data were used. Cell annotations for each dataset were based on publicly available data. The AddModuleScore function was performed using the Seurat package[38] to calculate the average expression levels in each protein community. The combined score was calculated by multiplying the expression level by the proportion of positive cells in each community.

### Cell type composition analysis based on human brain and microglia snRNA-seq data

For a cell type composition analysis, publicly available human microglia[21,22] snRNA-seq data were used. ANOVA was used to compare the frequencies of different microglial subtypes across various conditions (Control, Aβ+, and Aβ+Tau+). Next, for the microglial subtypes “Microglia_GPNMB_LPL” and “Microglia_GPNMB_PLAT,” a Tukey HSD test was used to determine specific differences between groups (for cases where ANOVA showed p < 0.05).

### Gene Ontology functional enrichment analysis

A Gene Ontology (GO) analysis was used to explore the biological functions of our targeted gene set. The GO biological process (GObp) dataset version 2023.1 was downloaded from the Molecular Signatures Database (MsigDB) (https://www.gsea-msigdb.org/gsea/msigdb). For the analysis, the enrichment function in the clusterProfiler package was used, and values of p < 0.05, after BH correction, were considered significant. The Rrvgo[39] software was used to summarize the GO terms.

### Modulators of the A**β**-tau interaction

To explore modulators of the Aβ-tau interaction based on proteomics data, a slightly modified version of the MINDy algorithm[40] was employed. Briefly, (1) Euclidean distances for APP and MAPT were calculated using BIONIC features, and common elements within the top 5% with respect to proximity were selected for analyses. (2) For each candidate modulator, the top and bottom 35% of proteins with respect to expression levels were segregated, and mutual information for APP and Tau was calculated for each group. Mutual information, which measures the interdependence between two random variables X and Y, was defined using equation (1) as follows:

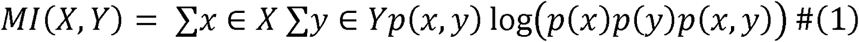

Here,

*p(x,y)* represents the joint probability distribution of *X* and *Y*.

*p(x)* and *p(y)* are the marginal probability distributions of *X* and *Y*, respectively. The logarithm base 2 was used.

Subsequently, (3) the difference in mutual information between the top and bottom 35% groups was calculated using equation (2):

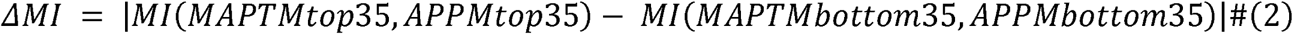

Here, M_top35_ and M_bottom35_ indicate the APP and MAPT groups in the top 35% and bottom 35% of protein expression levels, respectively. This differentiation is crucial for understanding the difference in mutual information between the two groups, allowing for a clear analysis of the Aβ-Tau interaction modulators. Bootstrapping with 1,000 iterations was then used to compare mutual information, with values of p < 0.05 deemed significant.

Next, factors moderating Aβ-Tau interactions were extracted using data obtained from BIONIC and a network analysis was conducted. Among candidate modulators identified in the MINDy analysis, HSPA5, which is closely associated with Aβ[41] and Tau[42], was included in the analysis. The inner product of the BIONIC-derived features was calculated to determine protein similarity, retaining the top 5% for network construction. From this network, the top 5% of factors based on Euclidean distances to *APP*, *MAPT*, and *HSPA5* were retained to form a subnetwork. Community detection within this subnetwork was performed using the leading eigenvector function of the iGraph package.[43]

### Neuropathological and RNA-seq datasets

The neuropathological and RNA-Seq datasets used in this study were obtained from previously published works.[44] Neuropathological data include quantitative immunostaining results for the microglial marker IBA1, astrocyte marker GFAP, Aβ, tau, and phosphorylated tau. The datasets included 377 samples collected from the cortical grey (parietal and temporal) and white matter (parietal) as well as the hippocampus. Additionally, the frequency distributions of age, sex, CERAD scores, and Braak scores among the participants were evaluated, differentiating between the no-dementia and dementia groups. Age was compared between the dementia and no-dementia groups using a t-test, whereas sex, Braak score, and CERAD score were compared between the groups using Fisher’s exact test.

### Linear regression analysis for exploring interaction terms

To explore moderating factors in the Aβ-tau interaction, a linear regression analysis was used to evaluate interaction effects, as described previously.[13] Measurements were performed on formalin-fixed, paraffin-embedded tissue sections (5 μm). For Aβ, the percentage of area covered by Aβ immunoreactivity was used, identifying both Aβ40 and Aβ42. The analysis focused on the percentage of the area covered by Tau2 immunoreactivity, identifying mature neurofibrillary tangles and dystrophic neurites.

AT8, an antibody that recognizes phospho-tau (Ser202, Thr205), was also utilized. Interaction effects with Aβ were investigated using immunoreactivities of IBA1 and GFAP and gene expression levels of *GPNMB*. IBA1 was measured as the percentage of area covered by IBA1 immunoreactivity, GFAP as the percentage of area covered by GFAP immunoreactivity (identifying activated astrocytes), and *GPNMB* gene expression levels were quantified as fragments per kilobase of exon per million reads mapped (FPKM). The interaction effects were evaluated using the *t*-test, with values of p < 0.05 considered significant.

### Trajectory analysis of human microglia snRNA-seq data

A trajectory analysis was conducted using human microglial snRNA-seq data and the Slingshot package. The default parameters for the slingshot function were utilized, and the getCurves function was used with the following parameters: approx_points = 300, threshold = 0.01, stretch = 0.8, allow.breaks = FALSE, and shrink = 0.99. From the dataset reported by Sun et al.,^21^ only microglia were extracted from immune cells for clustering. Clustering was performed using the Seurat package, following the tutorial provided by the Satija Lab (https://satijalab.org/seurat/articles/pbmc3k_tutorial). The settings were FindVariableFeatures with nfeatures = 4000, FindNeighbors, RunUMAP with dims = 1:10, and FindClusters with resolution = 0.5. To visualize gene expression over time, ggplot2 was used and regression curves were generated using generalized additive models (GAM).

### Heatmaps of CERAD and Braak scores in the ROSMAP cohort data

To investigate whether GPNMB protein expression moderates the Aβ-tau interaction in patients with MCI and early AD in the ROSMAP cohort, correlations between CERAD and Braak scores were evaluated. The CERAD score in the ROSMAP cohort is a semi-quantitative measure of neuritic plaque density, classified into four scores ranging from 4 to 1, corresponding to no AD, possible AD, probable AD, and definite AD, respectively. The Braak score is a semi-quantitative measure of the severity of neurofibrillary tangle pathology, ranging from 0 to 6, corresponding to Braak stages I–VI. For the top 35% and bottom 35% of patients based on GPNMB protein expression, the frequency of each CERAD and Braak score was visualized using a heatmap.

### Establishment of AlzPPMap

Using the R Shiny app, we developed a tool named AlzPPMap to analyze and visualize the integrated network. By inputting one or more gene names, the tool allows users to select a percentage of neighboring genes within the integrated network. The selected percentage of data is then used to construct and display a subnetwork.

### Results Study design

The analysis workflow, including an overview of the datasets, methods, and results, is presented in Supplementary Fig. S1. We extracted proteomics data for patients with MCI or early-stage AD, constructed a co-expression network, and integrated it with a PPI network using BIONIC, a deep learning-based method. To characterize the integrated data, we utilized enrichment analysis and whole snRNA-seq data (Fig. 2). Using proteomics data, we searched for factors associated with tau and Aβ (Fig. 3), constructed a subnetwork of tau, Aβ, and moderators, and performed cell type and functional enrichment analyses (Fig. 4). We validated these findings using immunohistochemical data from a separate cohort (Fig. 5). Additionally, for the subnetworks, we obtained microglia-specific snRNA-seq data to perform a microglial cell subtype enrichment analysis (Fig. 6). Finally, we validated these findings using immunohistochemical and gene expression data (Fig. 7).

**Fig. 2.**
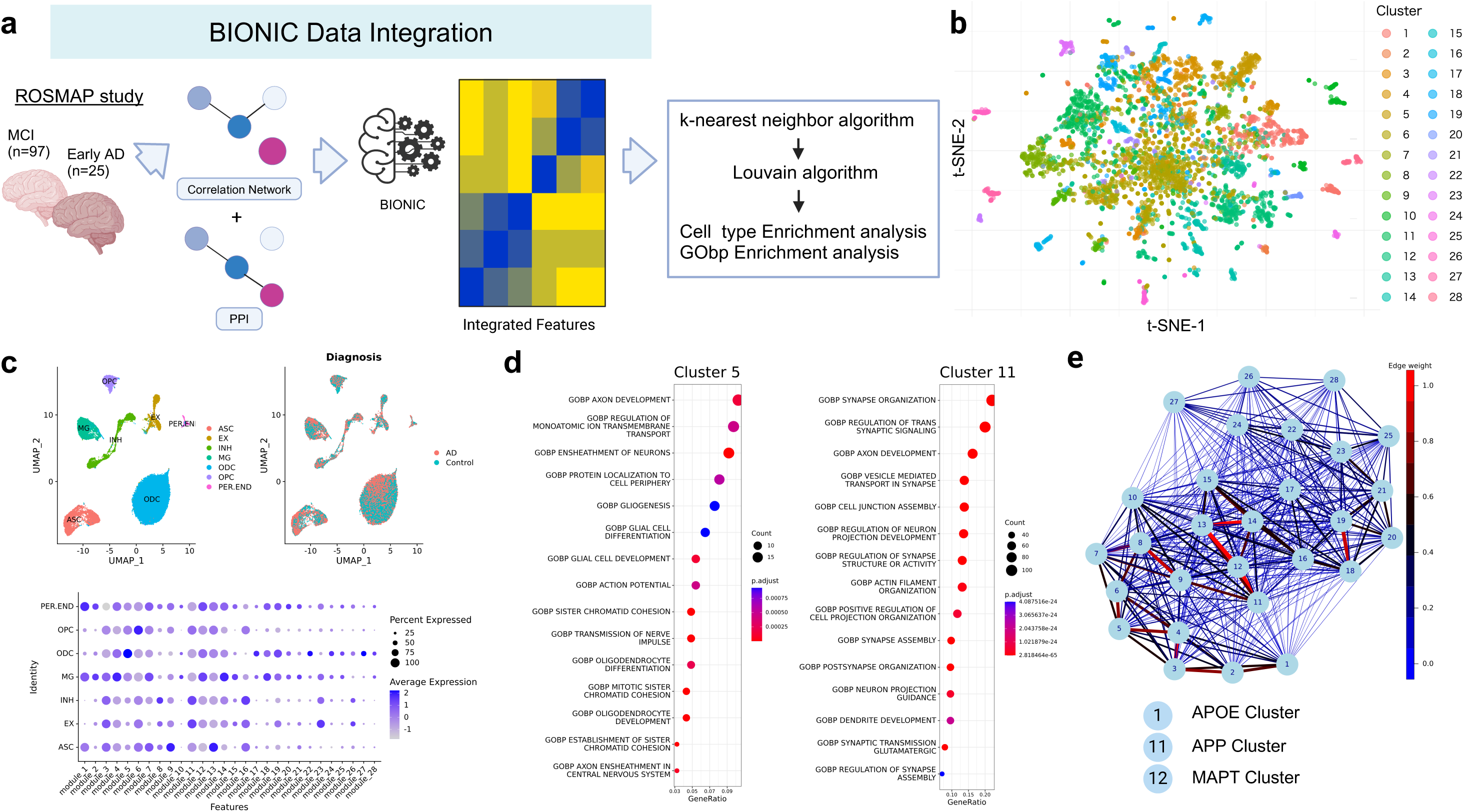
Characterization of integrated protein network modules. (a) Schematic representation of the workflow for integrating proteomics and protein–protein interaction (PPI) data and evaluation of integrated features. (b) Stochastic neighbor embedding (tSNE) plot displaying protein modules. (c) Upper panels: Uniform Manifold Approximation and Projection (UMAP) visualizations of single-nucleus RNA sequencing (snRNA-seq) data for patients with Alzheimer’s disease (AD) (GSE174367)[37] clustered according to cell types (left) and conditions (right) Lower panels: expression levels of proteins within the identified modules. (d) Representative Gene Ontology (GO) terms enriched in each protein module. (e) Network depicting the closeness among modules. Closeness was defined by averaging BIONIC-derived features within modules, calculating the Euclidean distances among modules, and taking reciprocals.

**Fig. 3.**
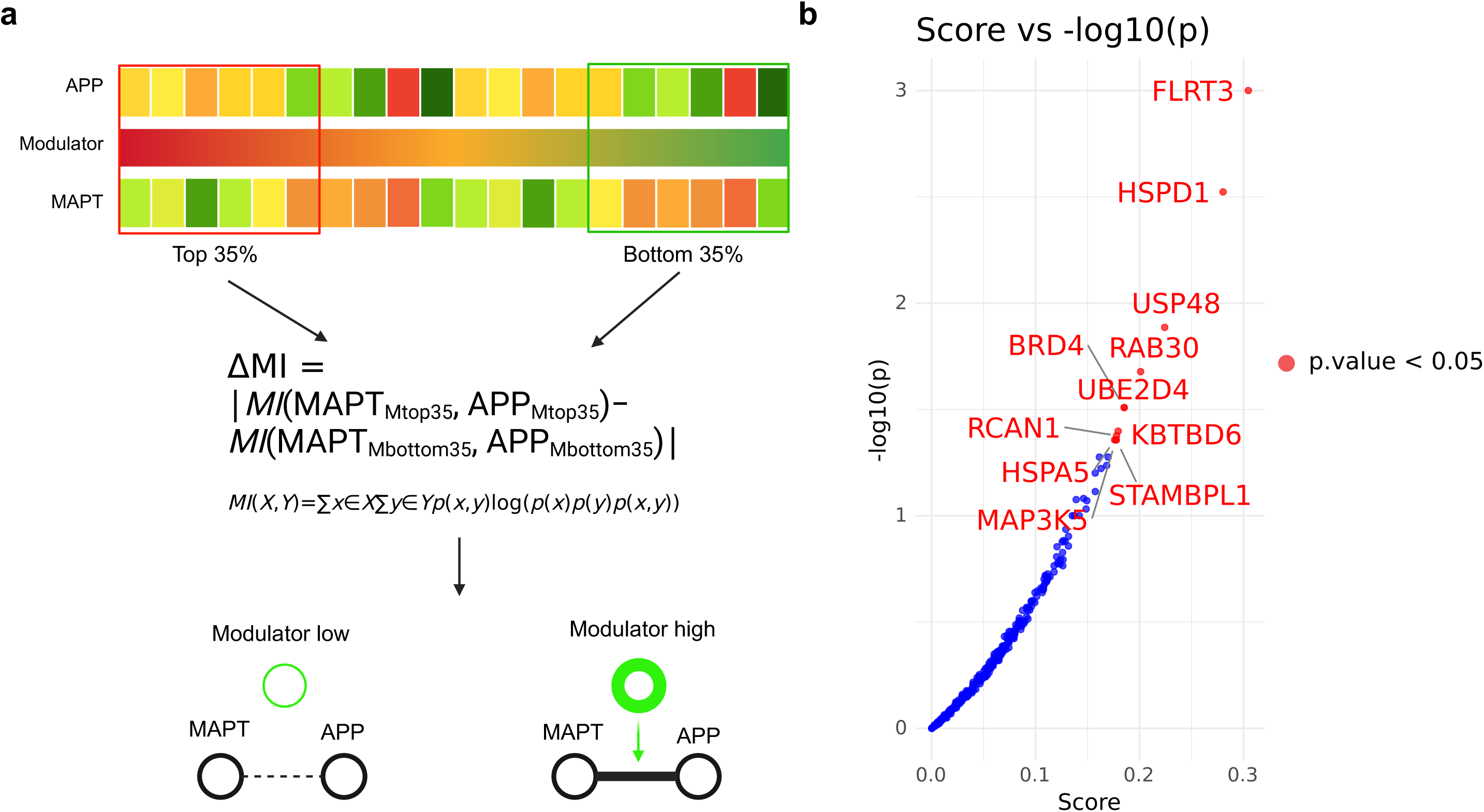
Inference of APP-MAPT modulators using a mutual information-based method. (a) Graphical representation of the Modulator Inference by Network Dynamics (MINDy) analysis. (b) Scatter plot illustrating the relationships between –log10(p-values) and Scores. Points representing p-values less than 0.05 are highlighted in red and are labeled with their corresponding gene symbols.

**Fig. 4.**
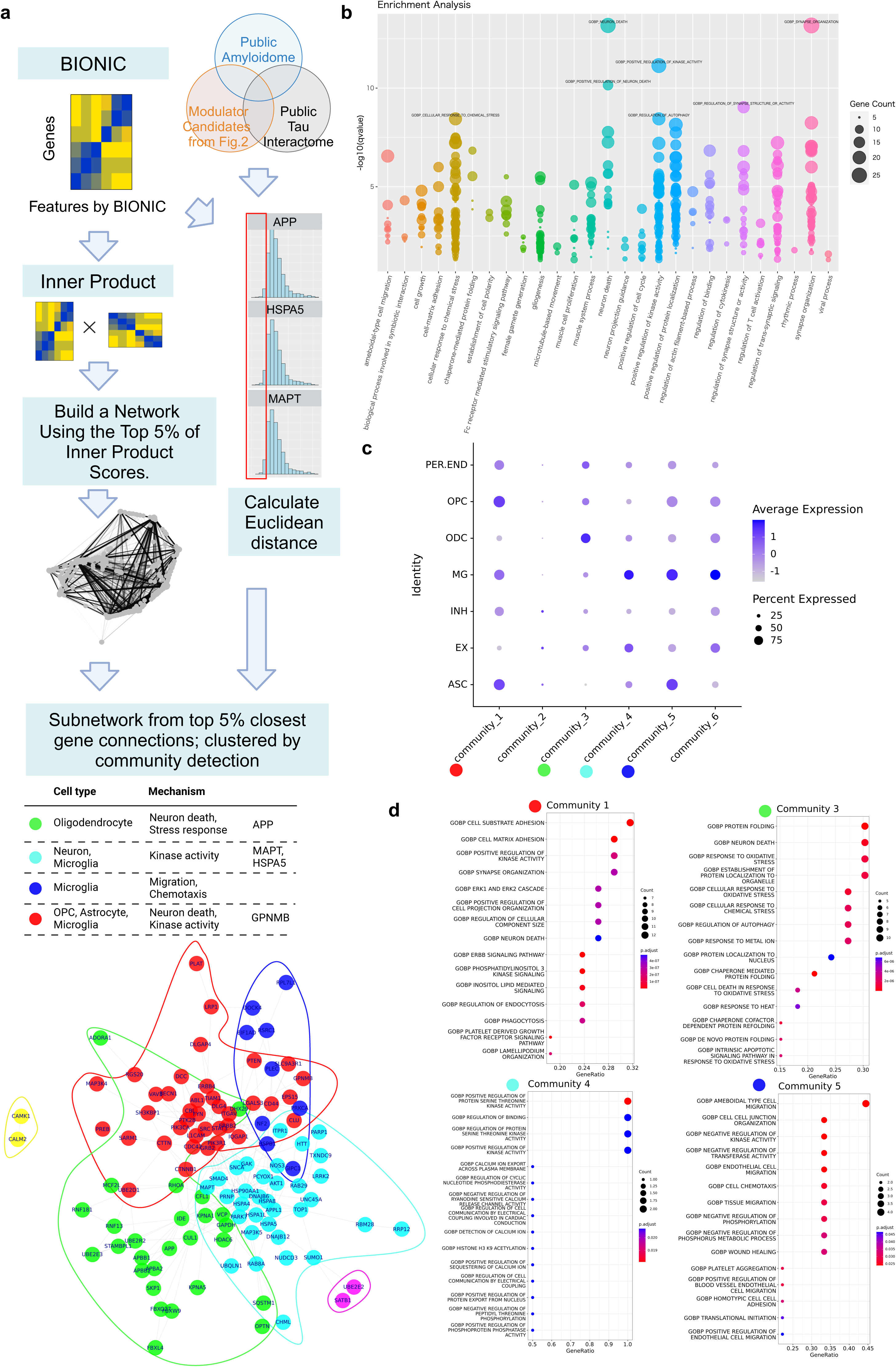
Glial cells moderate APP and MAPT interactions. (a) Workflow for network reconstruction from BIONIC analysis results, subnetwork construction using proteins highly associated with intermediary factors, APP and MAPT, and community detection within the network, followed by cell type identification and enrichment analyses for each community. (b) GO enrichment analysis of the subnetwork, categorized by parent terms using the rrvgo[39] software. (c) Representative GO terms enriched in each community. (d) Dotplot displaying expression levels of proteins within each community.

**Fig. 5.**
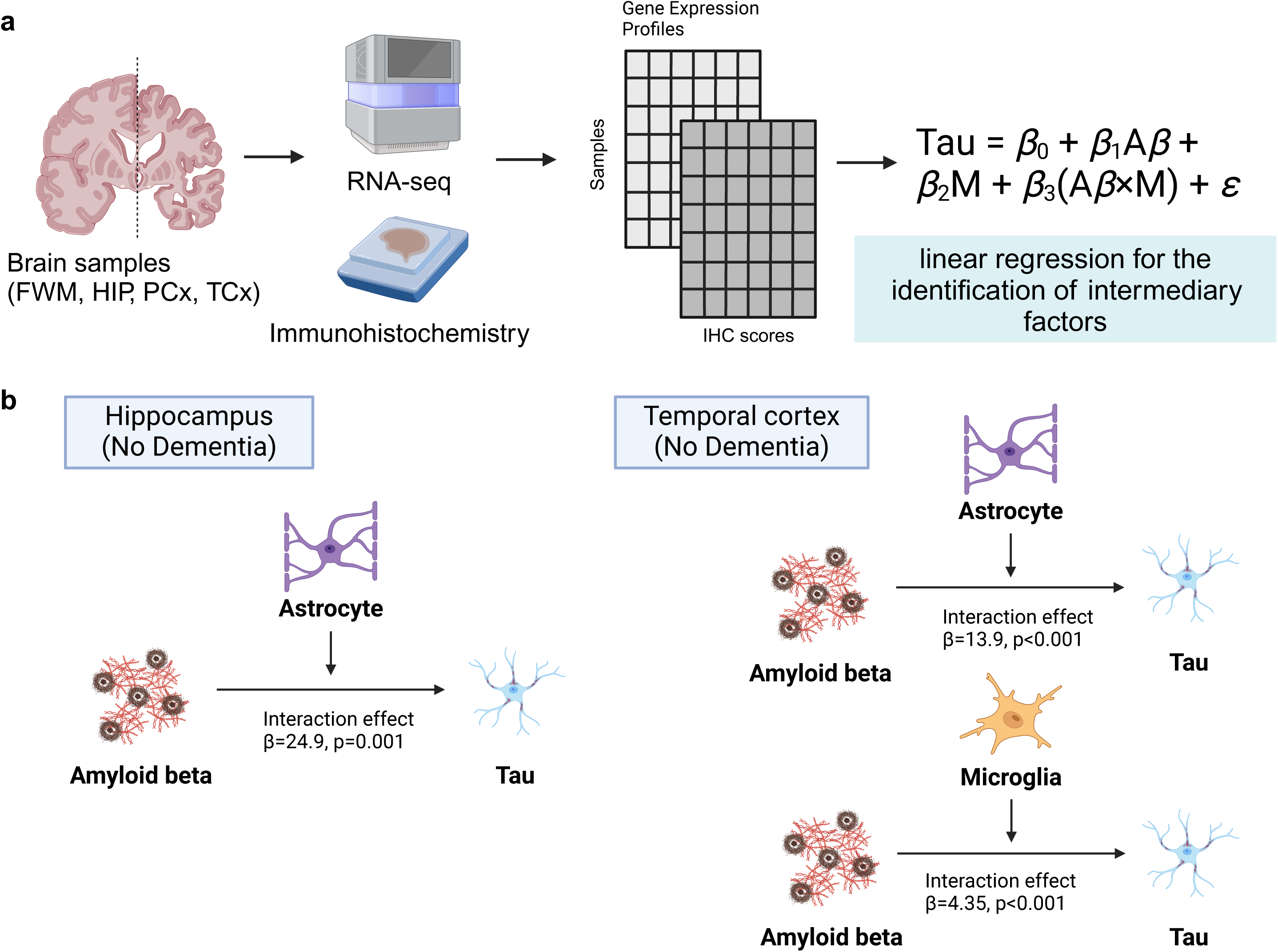
Histopathological analysis validated the roles of microglia/astrocytes as Aβ-tau moderators in early Alzheimer’s disease. (a) Schematic of the analytical workflow for linear regression analyses. Samples were sourced from four brain regions: frontal white matter (FWM), hippocampus (HIP), parietal neocortex (PCx), and temporal neocortex (TCx). RNA sequencing and immunohistochemistry data were used.[44] A linear regression was performed, repeated for GFAP and IBA1, and Aβ as predictors of tau interactions. (b) Significant results of linear regression analyses for each of the four distinct brain regions, with subjects categorized into no-dementia and dementia groups.

**Fig. 6.**
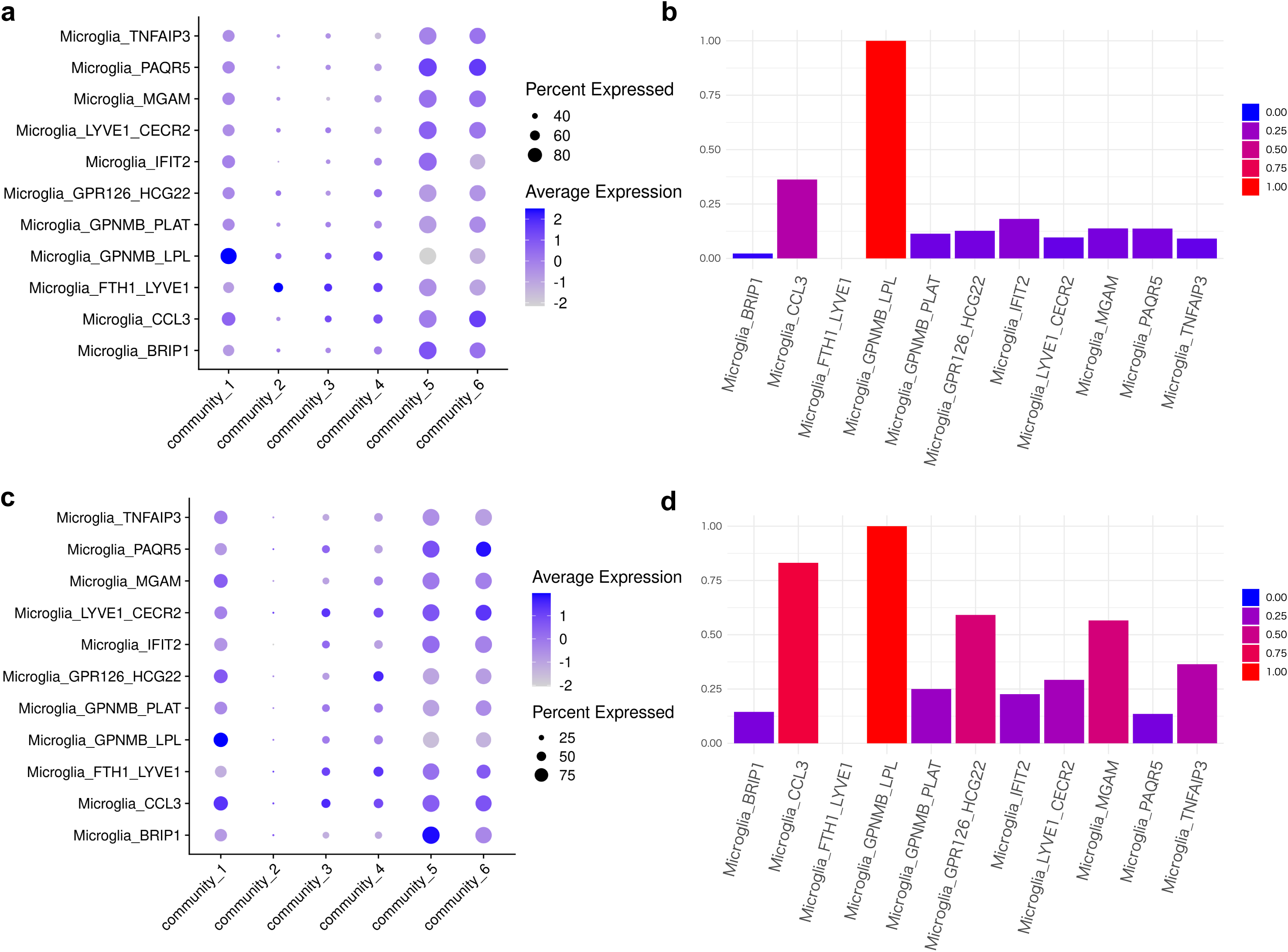
snRNA-seq Analysis Identifying Enrichment for GPNMB+ Microglia in Community 1. (a, c) Dot plot showing the expression levels of genes aggregated by community in each microglial subtype.[21,22] (b, d) Min-max normalized combined score for community 1, calculated by multiplying the expression level by the proportion of expressing cells.

**Fig. 7.**
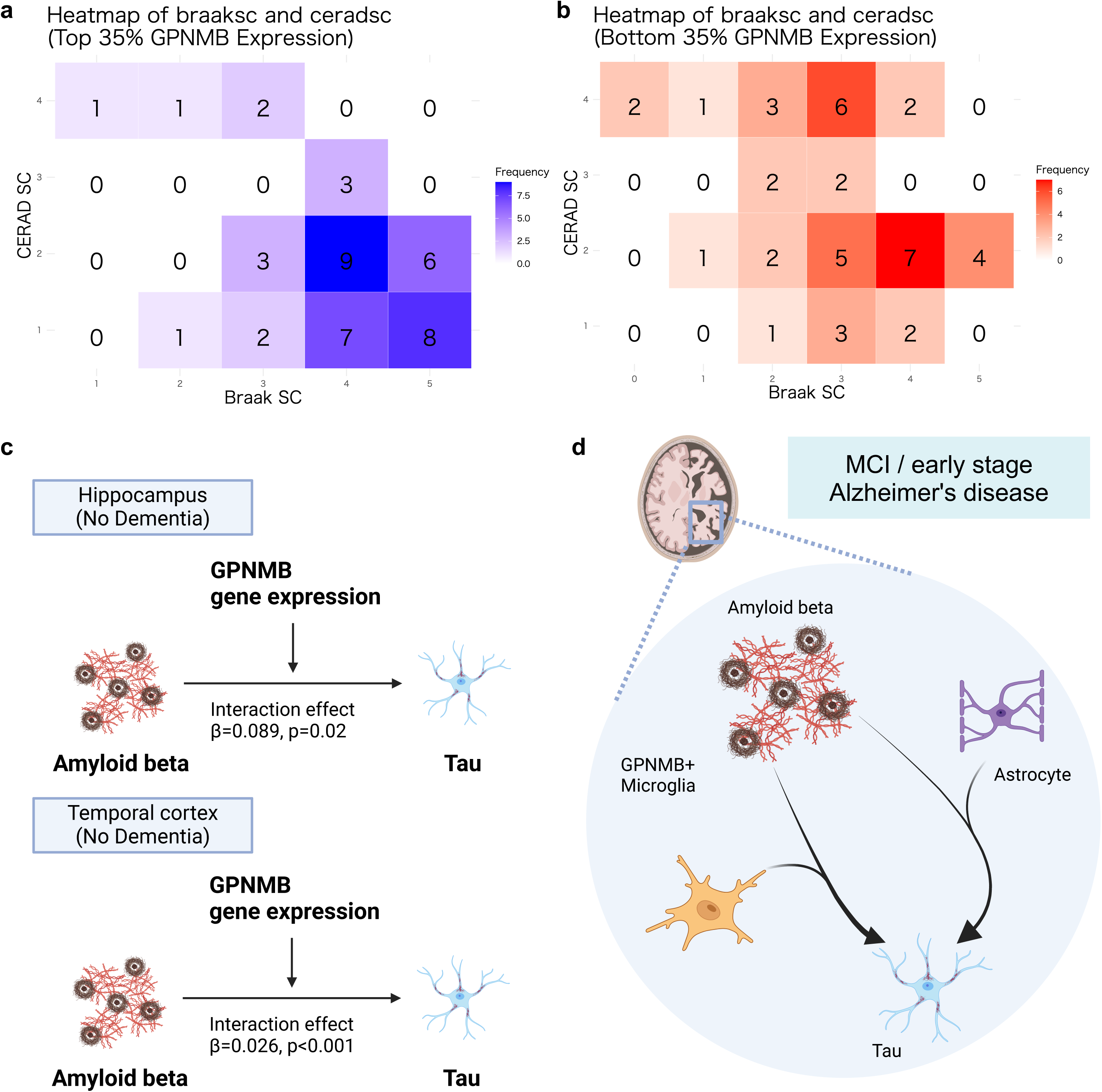
GPNMB+ microglia as a possible moderator of the. **A**β**-tau interaction** (a, b) Heatmaps displaying the correlations between CERAD and Braak scores in samples categorized by the top (a) and bottom (b) 35% of GPNMB expression. (c) Significant results of linear regression analyses for each of the four distinct brain regions, with subjects categorized into no-dementia and dementia groups. (d) Schematic representation summarizing the findings supporting the roles of both astrocytes and GPNMB+ microglia in the Aβ-tau interaction.

### Integration of protein co-expression and physical PPI networks

We constructed an integrated framework as a basis for analyzing the effects of PPIs and their moderators in the AD postmortem brain. To this end, we integrated proteomic data and a physical PPI network to construct a functional-physical PPI model. We employed BIONIC,[30] a deep learning method designed to integrate multiple biological networks while preserving their integrity (Fig. 2a). BIONIC uses a graph attention network[34] to integrate multiple networks, maintain the distinctive characteristics of each, and generate new features. It shows good performance[30] and has been used in analyses of gene interactions in cancer.[45]

First, a functional protein interaction network was constructed based on protein abundance data. We compiled postmortem proteome profiles for patients with MCI and early-stage AD from the dorsolateral prefrontal cortex, which were available from the Religious Order Study (ROS) and Rush Memory and Aging Project (MAP) cohort (https://adknowledgeportal.synapse.org/). Next, we generated a PPI network using the STRING database[33] (https://string-db.org/) and constructed subnetworks by prioritizing “high confidence” interactions. Using BIONIC, the integrated framework was embedded into a lower-dimensional vector space, and the embedded coordinates were obtained in a 512-dimensional space. Our downstream analysis used 512 integrated network features.

### Characterizing integrated protein network modules

Next, we characterized the biological context captured by the integrated network by evaluating the cell types and biological functions associated with the protein modules. To investigate these modules, we built k-nearest neighbor graphs of proteins based on the integrated network to obtain clusters (see Methods and Figs. 2a, b). The cell types and biological processes corresponding to each module were inferred using publicly available snRNA-seq data[37] and Gene Ontology terms. The modules in the integrated network contained cell type signatures (Fig. 2c) and were enriched in biological processes related to these cell types (Supplementary Table S1 and Fig. 2d). For example, module 1 was characterized by endothelial cells, microglia, and astrocytes, modules 5, 17, 22, and 27 were associated with oligodendrocytes, modules 9 and 13 with astrocytes, and module 11 with neurons (Fig. 2c). Consistent with the cell type results, in module 5, terms related to oligodendrocytes, such as “Ensheathment of neuron” and “Oligodendrocyte differentiation,” were identified, whereas module 11 was predominantly enriched in synapse-related terms (Fig. 2d).

### Identification of *APP* and *MAPT* as adjacent modules in the integrated network

We explored the inter-modular relationships within the integrated network by specifically examining modules that included *APP* and *MAPT*, which serve as markers for Aβ and tau, respectively. The *APP* gene was located in module 11 and *MAPT* was located in module 12. Additionally, *APOE*, which plays a significant role in the accumulation and aggregation of Aβ, was assigned to module 1. We then constructed a network of module–module interactions (Fig. 2e, and Supplementary Fig. S2).

Notably, these three modules were in close proximity in the network, with a strong correlation between the *APP* and *MAPT* modules (Pearson correlation coefficient = 0.7667, p < 0.001). Additionally, strong correlations were observed between the *APOE* module and both the *APP* (*r* = 0.6479, p < 0.001) and *MAPT* modules (*r* = 0.6289, p < 0.001). Furthermore, within the network, the *MAPT* module was closest to the *APP* module based on Euclidean distances; the *APP* module was second closest to the *MAPT* module. These module-to-module correlations in early-stage AD support the interactions between *APOE*, *APP*, and *MAPT*. [46,47]

Next, we evaluated whether similar findings could be obtained without network integration. Employing the STRING database with settings for high confidence and physical interactions alone, no interactions were detected between the *APP* and *MAPT* modules. Similarly, *APOE* did not interact with the other genes (Supplementary Fig. S3). When examining protein expression data separately, *APP* and *MAPT* did not show significant correlations (Pearson’s correlation coefficient = −0.0465, p = 0.61). These results underscore the importance of our integrated network approach in capturing the biological interactions between Aβ and tau, which are crucial for understanding early-stage AD development.

### Stress response factors moderate A**β**-tau interactions

Next, we aimed to identify factors that could moderate the Aβ-tau interaction based on the integrated network. For this purpose, we prioritized proteins with strong moderating effects on APP-MAPT interactions against proteins close to the *APP* and *MAPT* modules. We utilized the computational framework of Modulator Inference by Network Dynamics (MINDy)[40] (Fig. 3a). Briefly, MINDy calculates mutual information for interaction pairs (such as *APP* and *MAPT*), conditioned by the expression of the modulator, which reflects the modulating effect. In the present analysis, we extracted the neighboring proteins of the *APP* and *MAPT* modules in the integrated network (see Methods, “Modulators of the Aβ-tau interaction”). We then stratified the samples from the proteomics dataset into groups representing the top and bottom 35% of expression for each neighboring protein and evaluated differences in the magnitude of mutual information between *APP* and *MAPT* (Fig. 3a). We analyzed 312 neighboring proteins and identified the 11 most significant modulators (Fig. 3b). These candidates included factors involved in the cellular stress response (*HSPD1* and *HSPA5*),[48] ubiquitin-proteasome system (*USP48*, *KBTBD6*, *UBE2D4*, and *STAMBPL1*),[49–51] and apoptosis and cellular responses (*MAP3K5*).[51] These results suggest that proteostasis involving stress response genes is related to the Aβ-tau interaction.

To further identify proteins contributing to the Aβ-tau interaction from the moderator candidates, we investigated proteins known to physically interact with Aβ and/or tau sourced from the Amyloidome[41] and Tau Interactome.[42] Consequently, Heat Shock Protein Family A (Hsp70) Member 5 (*HSPA5*) was identified. *HSPA5*, a marker of ER stress associated with AD,[52,53] has been shown to interact in vitro with Aβ and tau, mitigating their toxicity[54–58]. Although in vivo studies have identified it as a candidate therapeutic target[59,60], the detailed mechanisms of action and its relevance in the Aβ-tau interaction are not fully understood[61].

### Glial cells are closely related to the APP-MAPT interaction in the integrated network

We further investigated whether stress responses involving *HSPA5* are involved in Aβ-tau interactions and identified related cell types and biological pathways. To address this, we exploited the integrated network extensively and reconstructed *HSPA5* and its neighboring subnetworks as well as *APP* and *MAPT* (Fig. 4a). Note that the subnetwork included other significant moderators, *STAMBPL1* and *MAP3K5*, suggesting that these factors, in addition to *HSPA5*, could be associated with the Aβ-tau interaction. This subnetwork was characterized by functions in neuronal death, positive regulation of kinase activity, cellular response to chemical stress, synapse organization, and regulation of autophagy, confirming its relation to stress response pathways (Fig. 4b, see also Supplementary Table S2).

We classified the subnetwork proteins into six communities based on topology and then estimated the cell types and biological functions associated with each community (Figs. 4c, d, and Supplementary Table S3). We found that glial cells were the primary cell type related to the stress response subnetwork (Fig. 4c). Additionally, the combined score for each community, calculated by multiplying the expression level by the proportion of cells, showed enrichment for glial cells (Supplementary Fig. S4a). Similar results were obtained using snRNA-seq data from two different cohorts[21,22] and different regions (Supplementary Figs. S4b, c). However, communities 2 and 6 were excluded from this analysis owing to the small number of genes included. In the APP-related Community 3, oligodendrocytes were enriched in protein folding, oxidative stress responses, neuronal death, and autophagy. Community 4, which contained *MAPT* and *HSPA5*, was characterized by microglia and excitatory neurons with enrichment in kinase activity. Other communities such as communities 1 and 5 were also enriched in glial cells (Figs. 4c, d). Microglia were detected as a feature of three of the four significantly enriched submodules (Fig. 4c).

### Histopathological data analysis validated the roles of microglia and astrocytes in the A**β**-tau interaction in early AD

To further investigate the effect of Aβ-tau interaction, we examined publicly available histopathological (ACT study) data.[44] This dataset contains extensive multimodal histopathological information, including Aβ and tau pathology. The distributions of age, sex, CERAD score, and Braak score in groups with and without dementia are shown in Supplementary Fig. S5. Among these parameters, CERAD and Braak scores showed significant differences between the groups (Fisher’s exact test, p = 0.00237 and p = 0.000468, respectively). In the no-dementia group, the average CERAD score was 1.476 and the average Braak score was 3.421. Braak stages I and II indicate that neurofibrillary tangles are confined mainly to the entorhinal region of the brain, whereas stages III and IV involve the limbic regions, such as the hippocampus.

We analyzed the effect of the interaction between tau and Aβ using a linear modeling approach. Tau2, an antibody for a mature neurofibrillary tangles and dystrophic neurites, was used as the dependent variable, whereas Aβ, along with GFAP or IBA1 (activated astrocyte or microglia markers, respectively), was used as the independent variable. We investigated the interaction between GFAP or IBA1 and Aβ, stratifying the analysis by the dementia status and across four brain regions (Fig. 5). For GFAP, a significant positive contribution to the interaction was observed in the hippocampus of both the dementia and no-dementia groups (HIP, β = 43.59, p = 0.022 and β = 24.852 and p = 0.001, respectively) and in the temporal neocortex of the no-dementia group (TCx, β = 13.873 and p < 0.001). In the case of IBA1, a positive contribution trend was observed in the HIP of the no-dementia group, though the relationship was not significant (β = 4.678 and p = 0.338), whereas a significant positive contribution was observed in the TCx (β = 4.352 and p < 0.001). Conversely, in the dementia group, a significant negative contribution was noted in the TCx (β = −11.124, p = 0.04; Fig. 5b and Supplementary Table S4). Analyses of AT8, an antibody that recognizes phospho-tau (Ser202 and Thr205) showed the same trend in the no-dementia group: GFAP (β = 33.912 and p = 0.235 in HIP; β = 62.732 and p < 0.001 in TCx) and IBA1 (β = 14.442 and p = 0.36 in HIP; β = 16.344 and p < 0.001 in TCx) (Supplementary Table S5). Of note, interactions were observed only in the HIP and TCx regions, which are vulnerable in patients with early-stage AD.[62]

The average CERAD and Braak scores in the no-dementia group mentioned earlier suggest that, although not exhibiting dementia, this group exhibited mild progression of AD pathology (Supplementary Figs. S5c, d). Collectively, these results support the hypothesis that astrocytes and microglia moderate the Aβ-tau interaction in early-stage AD.

### GPNMB+ microglia function as moderators of the A**β**-tau interaction

We further examined microglial subtypes that contribute to Aβ-tau interactions. Microglia-overexpressing neuroinflammation-related processes can be assigned to subtypes that exhibit different cellular states under different AD conditions.[63] We evaluated whether a certain subtype of microglia moderates the Aβ-tau interaction in early-stage AD. We accessed recently reported snRNA-seq data[22] from frontal cortex biopsies of 52 patients with hydrocephalus (NPH). This cohort included samples with confirmed amyloid and tau pathologies, in which AD was in an asymptomatic stage. Of the 52 living biopsies, Aβ plaques were identified in 19 subjects, Aβ plaques and phosphorylated tau pathology in eight subjects, and no pathology in the remaining 25 subjects; for subjects with Aβ pathology, the subsequent onset of AD was determined in longitudinal studies. In total, 892,828 high-quality single-nucleus profiles were included in analyses. Here, we focused on the microglial cell populations.

We performed an enrichment analysis of each microglial subtype for the response-related subnetwork communities (communities 1–6) based on the integrated network. GPNMB+ microglia were significantly enriched in community 1 (Fig. 6). The expression level and proportion of cells identified as the GPNMB+ microglial subtype was high for the community 1 signature score (Fig. 6a). In addition, we compared the compositions of cell subtypes across different statuses in the same cohort. The frequency of GPNMB+ microglia was significantly higher in Aβ+ and Aβ+Tau+ samples than in the control group (Supplementary Fig. S6 and Supplementary Table S6). The combined score for Community 1, calculated by multiplying the expression level and the proportion of cells with positive expression, further confirmed that the GPNMB+ microglial subtype was the highest ranked among the examined subtypes (Fig. 6b). Importantly, the GPNMB+ microglial subtype is expressed in the early stages of AD and expands as the disease progresses.[22]

To further validate the involvement of GPNMB+ microglia in the Aβ-tau interaction in early-stage AD, we used another large-scale snRNA-seq data set for microglia subtypes in AD,[21] including transcriptome data for 194,000 single-nuclear microglia and multiple AD pathological phenotypes in 443 human subjects. The cell subtypes were annotated based on a previous study.[22] We found the same trend towards enrichment for GPNMB+ in Module 1 (Figs. 6c, d). These results suggest that the stress response module moderating Aβ-tau interactions in early AD corresponds to GPNMB+ microglia.

### A**β**-tau stress response mediators are activated prior to GPMNB+ microglial activation

We further examined the relationship between stress response-related mediators of the Aβ-tau interaction and GPNMB+ microglia. Previous studies have reported that ER stress induces GPNMB expression in neurons but not in microglia.[64] We hypothesized that stress response mediators are involved in the transition of microglia from a homeostatic state to an activated GPNMB+ state in the human brain. To test this hypothesis, trajectory analysis of the two snRNA-seq datasets used in the previous analyses was performed using Slingshot.[65] Briefly, in this analysis, cells are embedded in the manifold space based on expression similarity between cells. This trajectory reflects the cellular state and can be treated as a “pseudo-time” axis. Multiple branched trajectories were identified. For each trajectory, we identified the GPNMB+ microglial coordinates on the pseudo-time trajectories (Supplementary Figs. S7a, c).

Next, we analyzed the points at which the candidate Aβ-tau-moderating factors related to the stress response predicted using MINDy were expressed in the pseudo-time analysis. In both datasets, the moderator proteins were highly expressed prior to GPNMB+ microglia in all pseudo-time trajectories (Supplementary Figs. S7b, d). Similar results were obtained for *HSPA5*, a moderating factor based on the amyloidome (i.e., the collection of amyloid peptides and related proteins, peptides, and other molecules)[41] and tau interactome.[42] These results suggest that stress factors moderating the Aβ-tau interaction may act through the activation of GPNMB+ microglia.

### *GPMNB* protein expression is correlated with the clinical severity of AD

We evaluated the hypothesis that GPMNB+ microglia moderate Aβ-tau interactions in early AD. In particular, we examined whether *GPMNB* could serve as a biomarker for the clinical severity of AD. For this purpose, we assessed brain tissue protein expression and clinical parameters from the ROSMAP cohort data using CERAD and Braak scores as indicators of the clinical severity of AD. We stratified the patients into two groups based on *GPNMB* protein expression: the top 35% and bottom 35%. The *GPNMB* high- and low-expression groups corresponded to the GPNMB+ microglia-activated and non-activated groups, respectively. Within the ROSMAP cohort, the CERAD score is a semi-quantitative measure of neuritic plaques, where 1 indicates definite AD, 2 is probable AD, 3 is possible AD, and 4 represents no AD. The top 35% showed greater disease progression based on both CERAD and Braak scores than that of the bottom 35% (Figs. 7a, b).

### *GPNMB* shows significant A**β**-tau interaction effects in the preclinical stage

We further examined whether *GPNMB* moderates the effect of the Aβ-tau interaction in early-stage AD. To this end, we assessed the histopathological and AD diagnostic information from the ACT study as well as gene expression data. We tested interaction effects of *GPNMB* gene expression on Aβ and tau by linear regression analyses (Fig. 7c, Supplementary Tables S7 and S8).

There were significant positive interaction effects of *GPNMB* gene expression on Aβ-tau in the non-dementia group in the HIP and TCx brain regions (β = 0.089, p = 0.02 and β = 0.026, p < 0.001 in HIP and TCx, respectively). Notably, no significant interaction effects were observed at the same sites in the dementia group. Collectively, these findings suggest that GPNMB+ microglia and astrocyte moderate the Aβ-tau interaction in the early stages of AD (Fig. 7d).

### Establishment of an integrated network analysis tool, AlzPPMap

Finally, we developed a tool named AlzPPMap (Alzheimer’s disease Proteomics PPI Integrated Network Map) for the analysis and visualization of integrated networks. This tool enables users to analyze the integrated functional network derived from proteomics data alongside physical networks derived from PPIs. Users can select one or more genes of interest and, as demonstrated in this study, choose a specific percentage of closely related genes within the network based on their proximity to the selected gene(s). The selected genes are then used to construct, visualize, and detect communities within a subnetwork. AlzPPMap facilitates the identification of molecules and molecular pathways that are both functionally and physically associated with the target genes. The tool can be accessed at: https://igcore.cloud/GerOmics/AlzPPMap.

## Discussion

The number of individuals with AD is expected to continue increasing,[66] making the development of effective treatments a pressing issue. As mentioned in the Introduction, whereas Aβ and tau proteins are known to contribute to AD pathology, the details of their involvement remain unclear.[1,2] The specific interactions between them have been reported,[4–7] but not the underlying mechanisms. In this study, we employed a deep learning-based network integration analysis to investigate the mechanisms of the Aβ-tau interaction in patients with early-stage AD. The study was conducted in four steps: (1) network integration, (2) identification of moderators and subnetwork construction, (3) validation analysis using histopathological data, and (4) development of a user-friendly web application.

In the first phase, we constructed an integrated functional-physical interaction network where two different data types—protein co-expression and physical PPIs—were combined into a single network using BIONIC.[30] Interestingly, enrichment analysis and snRNA-seq data showed that the integrated network captured relevant cell types and pathways. We also confirmed that Aβ and tau were present in neighboring modules in the integrated network. Notably, neither co-expression information nor physical interaction data alone could capture the Aβ-tau interaction, emphasizing the need to evaluate functional and physical interactions simultaneously. Co-expression network analyses are widely used in AD research,[23–25] but it is difficult to represent physical interactions, such as Aβ-tau interactions or the amyloidome. Therefore, an approach that integrates networks based on different types of biological information may be effective in elucidating the complex molecular pathogenesis of AD.

In the second phase, we used the MINDy[40] algorithm to search for factors that moderate the Aβ-tau interaction using proteomics expression data. We identified ER stress markers such as *HSPA5*, along with other stress response-related factors, including *HSPD1*, *USP48*, *KBTBD6*, *UBE2D4*, *STAMBPL1*, and *MAP3K5*, as potential moderators. Although ER stress responses have been frequently reported to be associated with AD,[52–60] their detailed mechanisms remain unclear. In this study, our findings suggested that stress response pathways were associated with the Aβ-tau interaction, indicating that elucidating how these pathways influence the progression of AD could be a focus for future research. In particular, understanding how ER stress responses are involved in the Aβ-tau interaction and transmitted through glial cells could contribute to the development of novel therapeutic strategies. We next identified molecules, cell types, and pathways involved in the Aβ-tau interaction by constructing subnetworks from the factors identified in the second phase, extracting genes closely related to the two proteins within the integrated network. The results suggested that glial cells, particularly GPNMB+ microglia, are involved in the Aβ-tau interaction.

In the third phase, we validated the findings using an independent dataset with histopathological data from the ACT study.[44] Through linear regression analysis, we examined the interaction between GFAP, IBA1, and GPNMB with Aβ in the hippocampus and temporal neocortex in the non-dementia group. Significant positive interactions were detected, further suggesting that astrocytes and GPNMB+ microglia are moderators of the Aβ-tau interaction in early-stage AD. GPNMB+microglia, a recently identified subtype, are elevated in the frontal white matter and cortex of patients with AD.[67] Additionally, *GPNMB* is expressed in activated microglia[68] and is associated with AD pathology.[22] Furthermore, the expression of *GPNMB* in microglia is correlated with recently reported subtypes, such as disease-associated microglia (DAM)[63] and microglial neurodegenerative phenotype (MGnD),[69] which are closely associated with plaques and neurodegeneration. However, their specific functions related to Aβ and tau have not been fully elucidated. Our results provide new insights into the functions of GPNMB+ microglia in the pathogenesis of AD.

In summary, network integration analysis has been extensively reported in cancer research,[26–30] but its use in AD research remains limited. Through this study, we were able to identify potential moderators of the Aβ-tau interaction by integrating proteomics and PPI data using BIONIC.[30] Future analyses using a framework like the one presented in this study, integrating other omics data, could lead to new insights into the mechanisms of AD and other disease areas. Additionally, in the last phase, we developed a tool named AlzPPMap to evaluate the integrated network constructed in this study. This tool enables the analysis and visualization of relationships between other genes, potentially providing new insights and ideas for experimental validation.

This study had several limitations. First, except for the GPNMB+ microglial data[22], all data were derived from postmortem brains. This introduces potential confounding factors between postmortem effects and AD pathology. Second, in the analyses summarized in Fig. 4, protein expression was aggregated by genes, not accounting for multiple isoforms or post-translational modifications, especially in *APP* and *MAPT*. For example, Aβ40 and Aβ42 are produced from *APP* through different processes, and Tau in *MAPT* undergoes various phosphorylation modifications,[70] impacting the disease differently. Histopathological evaluations were used to validate the integrated protein network analysis results. Third, the *GPNMB* gene expression analysis was based on bulk RNA-seq data, which may not specifically reflect GPNMB+ microglia. A detailed analysis of the role of GPNMB+ microglia in AD requires further experimental studies. Fourth, although we applied a significance threshold of P < 0.05 with the Benjamini-Hochberg correction, there is ongoing discussion in the field regarding the potential benefits of using a more stringent threshold. Future studies with larger sample sizes could help to further support our findings. Nonetheless, our findings offer valuable new insights into the Aβ-tau interaction and highlight potential directions for the development of AD therapeutics.

## List of abbreviations

AD: Alzheimer’s disease

Aβ: amyloid-beta

DAM: disease-associated microglia

MCI: mild cognitive impairment

PPI: protein–protein interaction

ROSMAP: Religious Order Study and Rush Memory and Aging Project

snRNA-seq: single-nucleus RNA sequencing

TMT: tandem mass tag

## Declarations Acknowledgments

We extend our thanks to all of the donors who generously contributed invaluable data for this study. We are also grateful to the researchers who made their data publicly available. The results published here are, in whole or in part, based on data obtained from the AD Knowledge Portal (https://adknowledgeportal.org). The study data were provided by the Rush Alzheimer’s Disease Center, Rush University Medical Center, Chicago. Data collection was supported through funding by NIA grants P30AG10161 (ROS), R01AG15819 (ROSMAP; genomics and RNAseq), R01AG17917 (MAP), R01AG30146, R01AG36042 (5hC methylation, ATACseq), RC2AG036547 (H3K9Ac), R01AG36836 (RNAseq), R01AG48015 (monocyte RNAseq) RF1AG57473 (single nucleus RNAseq), U01AG32984 (genomic and whole exome sequencing), U01AG46152 (ROSMAP AMP-AD, targeted proteomics), U01AG46161(TMT proteomics), U01AG61356 (whole genome sequencing, targeted proteomics, ROSMAP AMP-AD), the Illinois Department of Public Health (ROSMAP), and the Translational Genomics Research Institute (genomic). Additional phenotypic data were obtained from www.radc.rush.edu. Human AD brain single-nucleus RNA sequencing (snRNA-seq) data were downloaded from the NCBI for Biotechnology Information Gene Expression Omnibus (https://www.ncbi.nlm.nih.gov/geo/). Microglial snRNA-seq data were accessed at https://braincelldata.org/resource and https://doi.org/10.7303/syn2580853. Additionally, the human neuropathological and RNA sequencing datasets used for the linear regression analysis are available at NIAGADS (https://www.niagads.org/). Figures were created using Bio-Render.com.

## Author contributions

Conceptualization and Methodology: M.Y., K.A., Formal Analysis, Investigation, Data Curation, Writing – Original Draft, Visualization: K.A., Resource, Writing – Review & Editing, Supervision, Project Administration and Funding Acquisition: M.Y.

## Human Ethics and Consent to Participate declarations

This study utilizes publicly available datasets obtained through appropriate agreements or public data portals, ensuring compliance with ethical guidelines. The proteomic data used in this study were accessed from the AD Knowledge Portal (https://adknowledgeportal.org) under the terms of the data use agreement. These datasets were collected with the informed consent of participants and ethical approval from the original study institutions, including ROSMAP cohort studies. Additionally, neuropathological and single-nucleus RNA sequencing (snRNA-seq) datasets used in this study were obtained from the NCBI Gene Expression Omnibus (GEO) and NIAGADS (https://www.niagads.org/), where all samples were collected following approved institutional protocols, with participants providing informed consent for research and data sharing. For data requiring restricted access, the necessary approvals and permissions were obtained from the respective data providers. This study adhered to all relevant ethical regulations for the analysis of secondary human data.

## Availability of data and materials

All data used in this study are publicly accessible. The proteomic data can be accessed via the AD Knowledge Portal (https://doi.org/10.7303/syn2580853). Human AD brain snRNA-seq and microglia snRNA-seq datasets are available at the NCBI Gene Expression Omnibus (GEO) under accession number GSE174367 and via the following links: https://braincelldata.org/resource and https://doi.org/10.7303/syn2580853. The human neuropathological and RNA sequencing datasets employed for the linear regression analysis can be found at https://www.niagads.org/datasets/ng00059. Additional information required to reanalyze the data reported in this study is available from the lead author upon request.

## Code availability

The code used can be found on GitHub (https://github.com/matsui-lab/AlzPPMap/tree/main).

## References

1. d’Errico P, Meyer-Luehmann M. Mechanisms of Pathogenic Tau and Aβ Protein Spreading in Alzheimer’s Disease. Front Aging Neurosci [Internet]. 2020;12:265. Available from: 10.3389/fnagi.2020.00265

2. Knopman DS, Amieva H, Petersen RC, Chételat G, Holtzman DM, Hyman BT, et al. Alzheimer disease. Nature Reviews Disease Primers [Internet]. 2021 [cited 2023 Nov 24];7:1–21. Available from: https://www.nature.com/articles/s41572-021-00269-y

3. Congdon EE, Ji C, Tetlow AM, Jiang Y, Sigurdsson EM. Tau-targeting therapies for Alzheimer disease: current status and future directions. Nat Rev Neurol [Internet]. 2023;19:715–36. Available from: 10.1038/s41582-023-00883-2

4. Bloom GS. Amyloid-β and tau: the trigger and bullet in Alzheimer disease pathogenesis. JAMA Neurol [Internet]. 2014;71:505–8. Available from: 10.1001/jamaneurol.2013.5847

5. Busche MA, Hyman BT. Synergy between amyloid-β and tau in Alzheimer’s disease. Nat Neurosci [Internet]. 2020;23:1183–93. Available from: 10.1038/s41593-020-0687-6

6. Zhang H, Wei W, Zhao M, Ma L, Jiang X, Pei H, et al. Interaction between Aβ and Tau in the Pathogenesis of Alzheimer’s Disease. Int J Biol Sci [Internet]. 2021;17:2181–92. Available from: 10.7150/ijbs.57078

7. Lee WJ, Brown JA, Kim HR, La Joie R, Cho H, Lyoo CH, et al. Regional Aβ-tau interactions promote onset and acceleration of Alzheimer’s disease tau spreading. Neuron [Internet]. 2022;110:1932–1943.e5. Available from: 10.1016/j.neuron.2022.03.034

8. Aksman LM, Oxtoby NP, Scelsi MA, Wijeratne PA, Young AL, Alves IL, et al. A data-driven study of Alzheimer’s disease related amyloid and tau pathology progression. Brain [Internet]. 2023;146:4935–48. Available from: 10.1093/brain/awad232

9. Josephs KA, Tosakulwong N, Weigand SD, Graff-Radford J, Schwarz CG, Senjem ML, et al. Flortaucipir PET uncovers relationships between tau and amyloid-β in primary age-related tauopathy and Alzheimer’s disease. Sci Transl Med [Internet]. 2024;16:eado8076. Available from: https://www.science.org/doi/10.1126/scitranslmed.ado8076

10. van Dyck CH, Swanson CJ, Aisen P, Bateman RJ, Chen C, Gee M, et al. Lecanemab in Early Alzheimer’s Disease. N Engl J Med [Internet]. 2023;388:9–21. Available from: 10.1056/NEJMoa2212948

11. Bellaver B, Povala G, Ferreira PCL, Ferrari-Souza JP, Leffa DT, Lussier FZ, et al. Astrocyte reactivity influences amyloid-β effects on tau pathology in preclinical Alzheimer’s disease. Nat Med [Internet]. 2023;29:1775–81. Available from: 10.1038/s41591-023-02380-x

12. Kim T, Yi D, Byun MS, Ahn H, Jung JH, Kong N, et al. Synergistic interaction of high blood pressure and cerebral beta-amyloid on tau pathology. Alzheimers Res Ther [Internet]. 2022;14:193. Available from: 10.1186/s13195-022-01149-7

13. Coomans EM, van Westen D, Pichet Binette A, Strandberg O, Spotorno N, Serrano GE, et al. Interactions between vascular burden and amyloid-β pathology on trajectories of tau accumulation. Brain [Internet]. 2023; Available from: 10.1093/brain/awad317

14. Yang H-S, Teng L, Kang D, Menon V, Ge T, Finucane HK, et al. Cell-type-specific Alzheimer’s disease polygenic risk scores are associated with distinct disease processes in Alzheimer’s disease. Nat Commun [Internet]. 2023 [cited 2023 Dec 7];14:1–13. Available from: https://www.nature.com/articles/s41467-023-43132-2

15. Shahidehpour RK, Nelson PT, Katsumata Y, Bachstetter AD. Exploring the link between dystrophic microglia and the spread of Alzheimer’s neuropathology. Brain [Internet]. 2024 [cited 2024 Aug 18]; Available from: https://pubmed.ncbi.nlm.nih.gov/39101580/

16. Mathys H, Peng Z, Boix CA, Victor MB, Leary N, Babu S, et al. Single-cell atlas reveals correlates of high cognitive function, dementia, and resilience to Alzheimer’s disease pathology. Cell [Internet]. 2023;186:4365–4385.e27. Available from: 10.1016/j.cell.2023.08.039

17. Zhou Y, Song WM, Andhey PS, Swain A, Levy T, Miller KR, et al. Human and mouse single-nucleus transcriptomics reveal TREM2-dependent and TREM2-independent cellular responses in Alzheimer’s disease. Nat Med [Internet]. 2020;26:131–42. Available from: 10.1038/s41591-019-0695-9

18. Olah M, Menon V, Habib N, Taga MF, Ma Y, Yung CJ, et al. Single cell RNA sequencing of human microglia uncovers a subset associated with Alzheimer’s disease. Nat Commun [Internet]. 2020;11:6129. Available from: 10.1038/s41467-020-19737-2

19. Grubman A, Chew G, Ouyang JF, Sun G, Choo XY, McLean C, et al. A single-cell atlas of entorhinal cortex from individuals with Alzheimer’s disease reveals cell-type-specific gene expression regulation. Nat Neurosci [Internet]. 2019;22:2087–97. Available from: 10.1038/s41593-019-0539-4

20. Mathys H, Davila-Velderrain J, Peng Z, Gao F, Mohammadi S, Young JZ, et al. Single-cell transcriptomic analysis of Alzheimer’s disease. Nature [Internet]. 2019;570:332–7. Available from: 10.1038/s41586-019-1195-2

21. Sun N, Victor MB, Park YP, Xiong X, Scannail AN, Leary N, et al. Human microglial state dynamics in Alzheimer’s disease progression. Cell [Internet]. 2023;186:4386–4403.e29. Available from: 10.1016/j.cell.2023.08.037

22. Gazestani V, Kamath T, Nadaf NM, Dougalis A, Burris SJ, Rooney B, et al. Early Alzheimer’s disease pathology in human cortex involves transient cell states. Cell [Internet]. 2023;186:4438–4453.e23. Available from: 10.1016/j.cell.2023.08.005

23. Neff RA, Wang M, Vatansever S, Guo L, Ming C, Wang Q, et al. Molecular subtyping of Alzheimer’s disease using RNA sequencing data reveals novel mechanisms and targets. Science Advances [Internet]. 2021;7:eabb5398. Available from: https://www.science.org/doi/abs/10.1126/sciadv.abb5398

24. Johnson ECB, Carter EK, Dammer EB, Duong DM, Gerasimov ES, Liu Y, et al. Large-scale deep multi-layer analysis of Alzheimer’s disease brain reveals strong proteomic disease-related changes not observed at the RNA level. Nat Neurosci [Internet]. 2022 [cited 2023 Nov 26];25:213–25. Available from: https://www.nature.com/articles/s41593-021-00999-y

25. Bai B, Wang X, Li Y, Chen P-C, Yu K, Dey KK, et al. Deep Multilayer Brain Proteomics Identifies Molecular Networks in Alzheimer’s Disease Progression. Neuron [Internet]. 2020;105:975–991.e7. Available from: 10.1016/j.neuron.2019.12.015

26. Cho H, Berger B, Peng J. Compact Integration of Multi-Network Topology for Functional Analysis of Genes. Cell Syst [Internet]. 2016;3:540–548.e5. Available from: 10.1016/j.cels.2016.10.017

27. Chen G, Liu Z-P. Graph attention network for link prediction of gene regulations from single-cell RNA-sequencing data. Bioinformatics [Internet]. 2022 [cited 2023 Dec 4];38:4522–9. Available from: https://academic.oup.com/bioinformatics/article/38/19/4522/6663989

28. Schulte-Sasse R, Budach S, Hnisz D, Marsico A. Integration of multiomics data with graph convolutional networks to identify new cancer genes and their associated molecular mechanisms. Nature Machine Intelligence [Internet]. 2021 [cited 2023 Dec 4];3:513–26. Available from: https://www.nature.com/articles/s42256-021-00325-y

29. Peng W, Tang Q, Dai W, Chen T. Improving cancer driver gene identification using multi-task learning on graph convolutional network. Brief Bioinform [Internet]. 2022;23. Available from: 10.1093/bib/bbab432

30. Forster DT, Li SC, Yashiroda Y, Yoshimura M, Li Z, Isuhuaylas LAV, et al. BIONIC: biological network integration using convolutions. Nat Methods [Internet]. 2022;19:1250–61. Available from: 10.1038/s41592-022-01616-x

31. ROSMAP_Proteomics TMT_BatchCorrectionOC.pdf.

32. Perneczky R, Wagenpfeil S, Komossa K, Grimmer T, Diehl J, Kurz A. Mapping scores onto stages: mini-mental state examination and clinical dementia rating. Am J Geriatr Psychiatry [Internet]. 2006;14:139–44. Available from: 10.1097/01.JGP.0000192478.82189.a8

33. Szklarczyk D, Gable AL, Lyon D, Junge A, Wyder S, Huerta-Cepas J, et al. STRING v11: protein-protein association networks with increased coverage, supporting functional discovery in genome-wide experimental datasets. Nucleic Acids Res [Internet]. 2019;47:D607–13. Available from: 10.1093/nar/gky1131

34. Veličković P, Cucurull G, Casanova A, Romero A, Liò P, Bengio Y. Graph Attention Networks [Internet]. arXiv [stat.ML]. 2017. Available from: http://arxiv.org/abs/1710.10903

35. Macosko EZ, Basu A, Satija R, Nemesh J, Shekhar K, Goldman M, et al. Highly Parallel Genome-wide Expression Profiling of Individual Cells Using Nanoliter Droplets. Cell [Internet]. 2015;161:1202–14. Available from: 10.1016/j.cell.2015.05.002

36. Blondel VD, Guillaume J-L, Lambiotte R, Lefebvre E. Fast unfolding of communities in large networks. J Stat Mech [Internet]. 2008 [cited 2024 Jun 8];2008:P10008. Available from: https://iopscience.iop.org/article/10.1088/1742-5468/2008/10/P10008/meta

37. Morabito S, Miyoshi E, Michael N, Shahin S, Martini AC, Head E, et al. Single-nucleus chromatin accessibility and transcriptomic characterization of Alzheimer’s disease. Nat Genet [Internet]. 2021;53:1143–55. Available from: 10.1038/s41588-021-00894-z

38. Hao Y, Hao S, Andersen-Nissen E, Mauck WM 3rd, Zheng S, Butler A, et al. Integrated analysis of multimodal single-cell data. Cell [Internet]. 2021;184:3573–3587.e29. Available from: 10.1016/j.cell.2021.04.048

39. Sayols S. rrvgo: a Bioconductor package for interpreting lists of Gene Ontology terms. MicroPubl Biol [Internet]. 2023;2023. Available from: 10.17912/micropub.biology.000811

40. Wang K, Saito M, Bisikirska BC, Alvarez MJ, Lim WK, Rajbhandari P, et al. Genome-wide identification of post-translational modulators of transcription factor activity in human B cells. Nat Biotechnol [Internet]. 2009;27:829–39. Available from: 10.1038/nbt.1563

41. Andrew RJ, Fisher K, Heesom KJ, Kellett KAB, Hooper NM. Quantitative interaction proteomics reveals differences in the interactomes of amyloid precursor protein isoforms. J Neurochem [Internet]. 2019;149:399–412. Available from: 10.1111/jnc.14666

42. Drummond E, Pires G, MacMurray C, Askenazi M, Nayak S, Bourdon M, et al. Phosphorylated tau interactome in the human Alzheimer’s disease brain. Brain [Internet]. 2020;143:2803–17. Available from: 10.1093/brain/awaa223

43. Csárdi G, Nepusz T. The igraph software package for complex network research. https://www.semanticscholar.org <..https://www.semanticscholar.org>. [Internet]. 2006 [cited 2024 Jun 8]; Available from: https://www.semanticscholar.org/paper/1d2744b83519657f5f2610698a8ddd177ced4f5c

44. Miller JA, Guillozet-Bongaarts A, Gibbons LE, Postupna N, Renz A, Beller AE, et al. Neuropathological and transcriptomic characteristics of the aged brain. Elife [Internet]. 2017;6. Available from: 10.7554/eLife.31126

45. Li P, Jiang Z, Liu T, Liu X, Qiao H, Yao X. Improving drug response prediction via integrating gene relationships with deep learning. Brief Bioinform [Internet]. 2024;25. Available from: 10.1093/bib/bbae153

46. Young CB, Johns E, Kennedy G, Belloy ME, Insel PS, Greicius MD, et al. APOE effects on regional tau in preclinical Alzheimer’s disease. Mol Neurodegener [Internet]. 2023;18:1. Available from: 10.1186/s13024-022-00590-4

47. Verghese PB, Castellano JM, Garai K, Wang Y, Jiang H, Shah A, et al. ApoE influences amyloid-β (Aβ) clearance despite minimal apoE/Aβ association in physiological conditions. Proc Natl Acad Sci U S A [Internet]. 2013;110:E1807–16. Available from: 10.1073/pnas.1220484110

48. Stetler RA, Gan Y, Zhang W, Liou AK, Gao Y, Cao G, et al. Heat shock proteins: cellular and molecular mechanisms in the central nervous system. Prog Neurobiol [Internet]. 2010;92:184–211. Available from: 10.1016/j.pneurobio.2010.05.002

49. Das S, Ramakrishna S, Kim K-S. Critical Roles of Deubiquitinating Enzymes in the Nervous System and Neurodegenerative Disorders. Mol Cells [Internet]. 2020;43:203–14. Available from: 10.14348/molcells.2020.2289

50. Genau HM, Huber J, Baschieri F, Akutsu M, Dötsch V, Farhan H, et al. CUL3-KBTBD6/KBTBD7 ubiquitin ligase cooperates with GABARAP proteins to spatially restrict TIAM1-RAC1 signaling. Mol Cell [Internet]. 2015;57:995–1010. Available from: 10.1016/j.molcel.2014.12.040

51. Ichijo H, Nishida E, Irie K, ten Dijke P, Saitoh M, Moriguchi T, et al. Induction of apoptosis by ASK1, a mammalian MAPKKK that activates SAPK/JNK and p38 signaling pathways. Science [Internet]. 1997;275:90–4. Available from: 10.1126/science.275.5296.90

52. Ajoolabady A, Lindholm D, Ren J, Pratico D. ER stress and UPR in Alzheimer’s disease: mechanisms, pathogenesis, treatments. Cell Death Dis [Internet]. 2022 [cited 2023 Dec 7];13:1–15. Available from: https://www.nature.com/articles/s41419-022-05153-5

53. Gorbatyuk MS, Gorbatyuk OS. The Molecular Chaperone GRP78/BiP as a Therapeutic Target for Neurodegenerative Disorders: A Mini Review. J Genet Syndr Gene Ther [Internet]. 2013;4. Available from: 10.4172/2157-7412.1000128

54. Yang Y, Turner RS, Gaut JR. The chaperone BiP/GRP78 binds to amyloid precursor protein and decreases Abeta40 and Abeta42 secretion. J Biol Chem [Internet]. 1998;273:25552–5. Available from: 10.1074/jbc.273.40.25552

55. Hoshino T, Nakaya T, Araki W, Suzuki K, Suzuki T, Mizushima T. Endoplasmic reticulum chaperones inhibit the production of amyloid-beta peptides. Biochem J [Internet]. 2007;402:581–9. Available from: 10.1042/BJ20061318

56. Sakono M, Kidani T. ATP-independent inhibition of amyloid beta fibrillation by the endoplasmic reticulum resident molecular chaperone GRP78. Biochem Biophys Res Commun [Internet]. 2017;493:500–3. Available from: 10.1016/j.bbrc.2017.08.162

57. Fontaine SN, Rauch JN, Nordhues BA, Assimon VA, Stothert AR, Jinwal UK, et al. Isoform-selective Genetic Inhibition of Constitutive Cytosolic Hsp70 Activity Promotes Client Tau Degradation Using an Altered Co-chaperone Complement. J Biol Chem [Internet]. 2015;290:13115–27. Available from: 10.1074/jbc.M115.637595

58. Moll A, Ramirez LM, Ninov M, Schwarz J, Urlaub H, Zweckstetter M. Hsp multichaperone complex buffers pathologically modified Tau. Nat Commun [Internet]. 2022 [cited 2023 Dec 7];13:1–13. Available from: https://www.nature.com/articles/s41467-022-31396-z

59. Shao Y, Xu Y, Di H, Shi X, Wang Y, Liu H, et al. The inhibition of ORMDL3 prevents Alzheimer’s disease through ferroptosis by PERK/ATF4/HSPA5 pathway. IET Nanobiotechnol [Internet]. 2023;17:182–96. Available from: 10.1049/nbt2.12113

60. Gao H, Lei X, Ye S, Ye T, Hua R, Wang G, et al. Genistein attenuates memory impairment in Alzheimer’s disease via ERS-mediated apoptotic pathway in vivo and in vitro. J Nutr Biochem [Internet]. 2022;109:109118. Available from: 10.1016/j.jnutbio.2022.109118

61. Castanho I, Murray TK, Hannon E, Jeffries A, Walker E, Laing E, et al. Transcriptional Signatures of Tau and Amyloid Neuropathology. Cell Rep [Internet]. 2020;30:2040–2054.e5. Available from: 10.1016/j.celrep.2020.01.063

62. Braak H, Braak E. Neuropathological stageing of Alzheimer-related changes. Acta Neuropathol [Internet]. 1991;82:239–59. Available from: 10.1007/BF00308809

63. Keren-Shaul H, Spinrad A, Weiner A, Matcovitch-Natan O, Dvir-Szternfeld R, Ulland TK, et al. A Unique Microglia Type Associated with Restricting Development of Alzheimer’s Disease. Cell [Internet]. 2017;169:1276–1290.e17. Available from: 10.1016/j.cell.2017.05.018

64. Noda Y, Tsuruma K, Takata M, Ishisaka M, Tanaka H, Nakano Y, et al. GPNMB Induces BiP Expression by Enhancing Splicing of BiP Pre-mRNA during the Endoplasmic Reticulum Stress Response. Sci Rep [Internet]. 2017;7:12160. Available from: 10.1038/s41598-017-11828-3

65. Street K, Risso D, Fletcher RB, Das D, Ngai J, Yosef N, et al. Slingshot: cell lineage and pseudotime inference for single-cell transcriptomics. BMC Genomics [Internet]. 2018;19:477. Available from: 10.1186/s12864-018-4772-0

66. 2024 Alzheimer’s disease facts and figures. Alzheimers Dement [Internet]. 2024;20:3708–821. Available from: https://pubmed.ncbi.nlm.nih.gov/38689398/

67. Satoh J-I, Kino Y, Yanaizu M, Ishida T, Saito Y. Microglia express GPNMB in the brains of Alzheimer’s disease and Nasu-Hakola disease. Intractable Rare Dis Res [Internet]. 2019;8:120–8. Available from: 10.5582/irdr.2019.01049

68. Hüttenrauch M, Ogorek I, Klafki H, Otto M, Stadelmann C, Weggen S, et al. Glycoprotein NMB: a novel Alzheimer’s disease associated marker expressed in a subset of activated microglia. Acta Neuropathol Commun [Internet]. 2018;6:108. Available from: 10.1186/s40478-018-0612-3

69. Krasemann S, Madore C, Cialic R, Baufeld C, Calcagno N, El Fatimy R, et al. The TREM2-APOE Pathway Drives the Transcriptional Phenotype of Dysfunctional Microglia in Neurodegenerative Diseases. Immunity [Internet]. 2017;47:566–581.e9. Available from: 10.1016/j.immuni.2017.08.008

70. Wegmann S, Biernat J, Mandelkow E. A current view on Tau protein phosphorylation in Alzheimer’s disease. Curr Opin Neurobiol [Internet]. 2021;69:131–8. Available from: 10.1016/j.conb.2021.03.003

