## Supplementary Information for "Integrative Network Analysis Reveals Novel Moderators of Aβ-Tau Interaction in Alzheimer’s Disease"

Yusuke Matsui

#### **Supplementary Fig. S1 Study Design**

A schematic workflow of how the dataset was filtered and utilized in each figure is depicted. Data used for analyses are outlined with black borders, while analyses are framed in blue borders and filled in blue. Specific data applied for sub-objectives are highlighted by black frames within the blue sections. The analysis was conducted in two phases: exploration and validation.

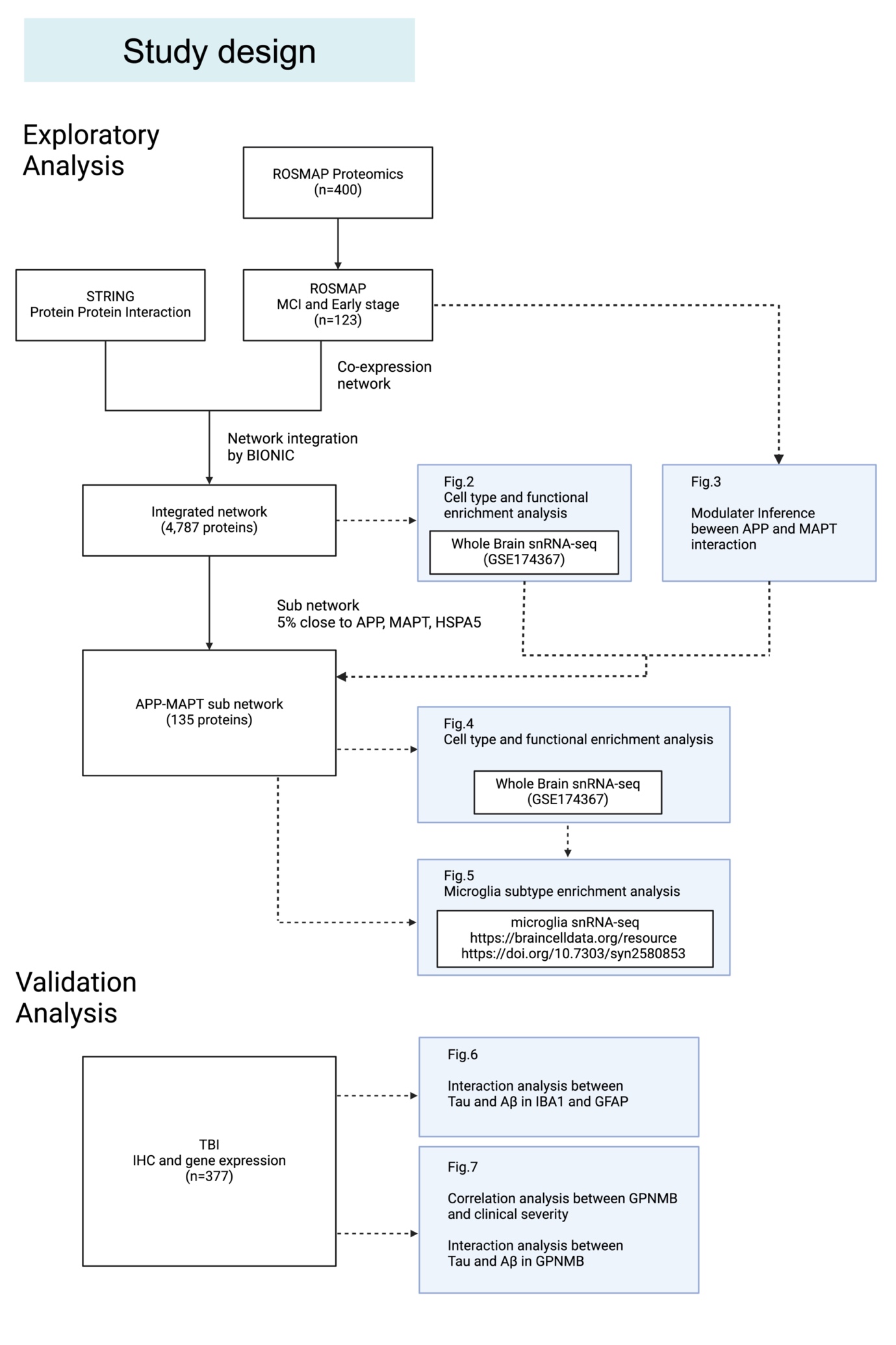

### **Supplementary Fig. S2. Distances and correlations among protein clusters, related to Fig. 3**

(a) Heatmap displaying the Euclidean distances between protein communities.

(b) Correlation plot representing the Pearson correlation coefficients among the identified protein communities.

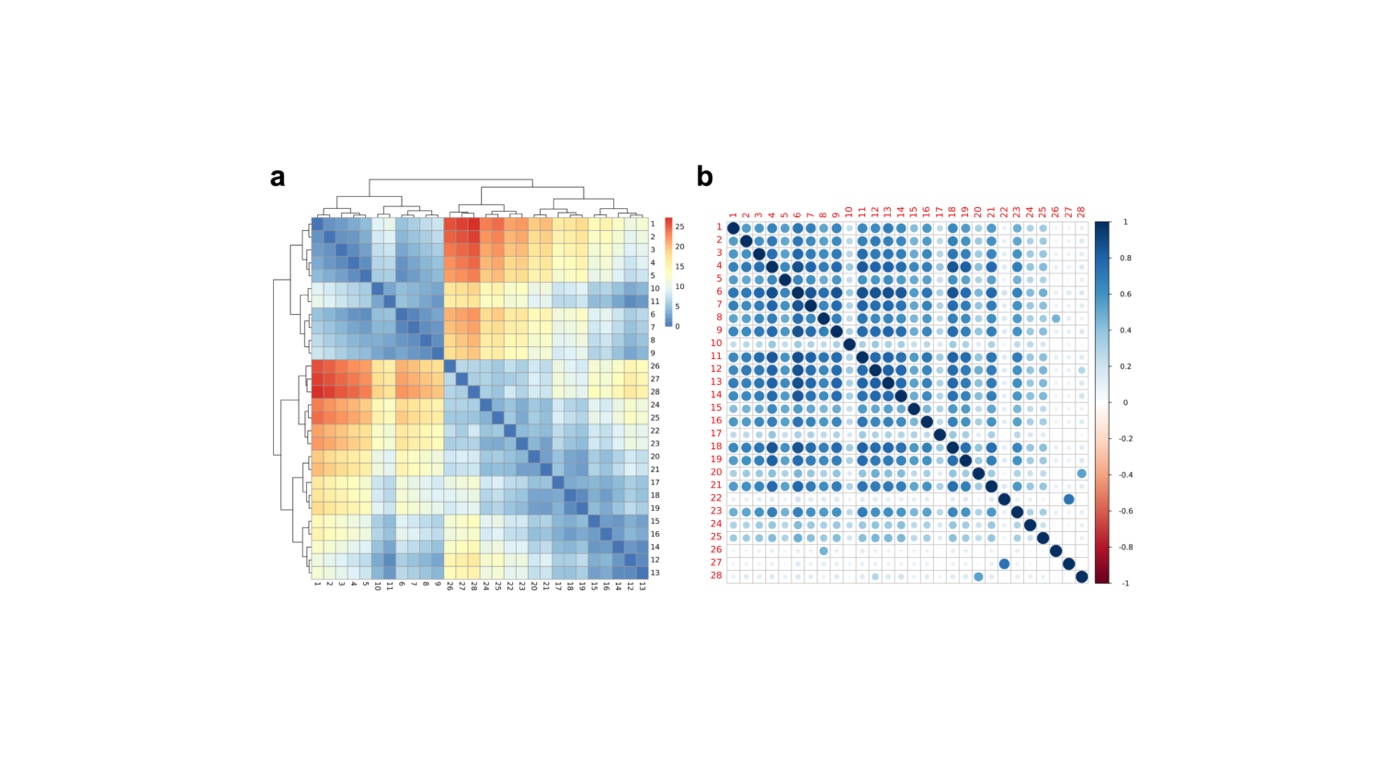

### **Supplementary Fig. S3. STRING Protein–Protein Interaction (PPI) Networks for MAPT and APP, related to Fig. 3**

Network diagrams displaying the PPI networks from STRING for (a) MAPT and (b) APP. In both networks, only interactions with a confidence score greater than 0.7, involving physical interactions and experimental validation, were included.

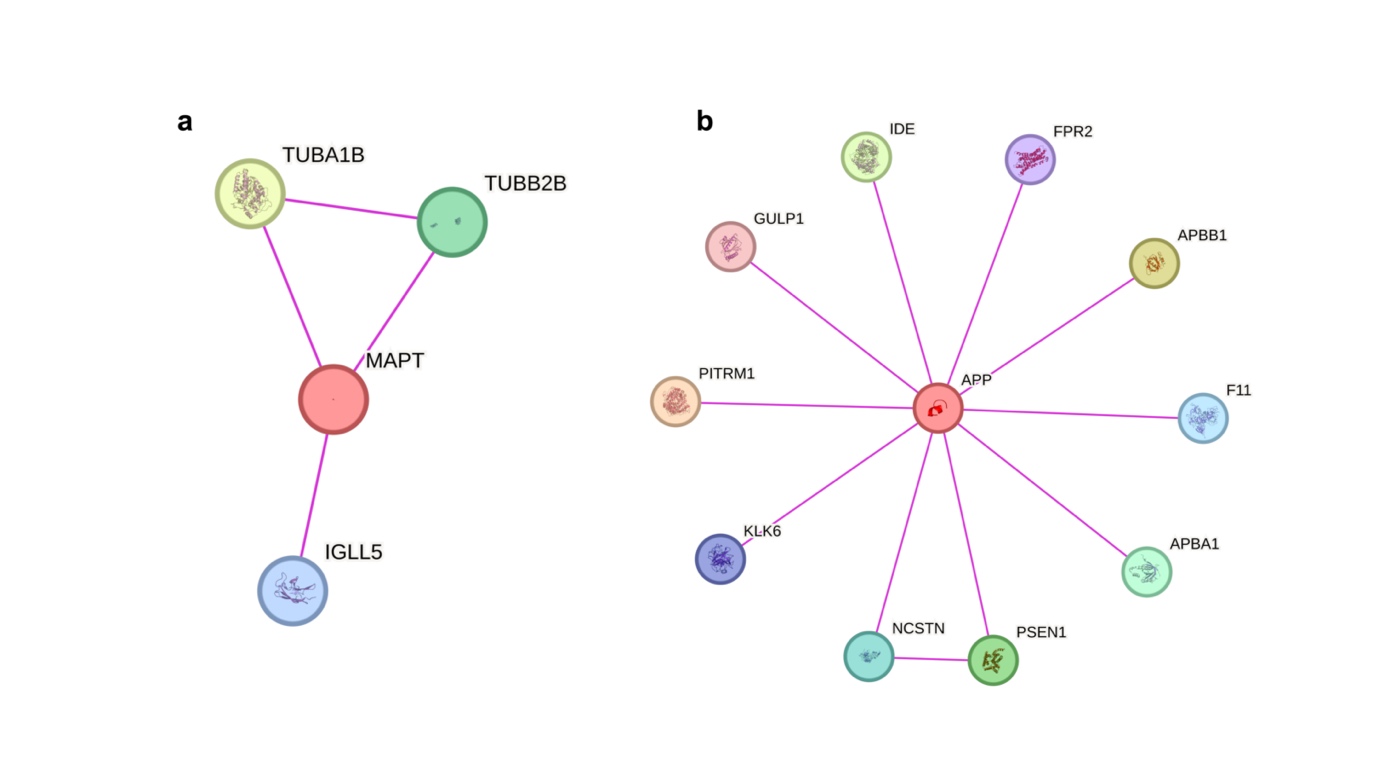

### **Supplementary Fig. S4 snRNA-seq Analysis Identifying Enrichment for Glial Cell Types in Subnetwork Communities, related to Fig. 4**

The min-max-normalized combined score for each community was calculated by multiplying the expression level by the proportion of expressing cells, based on the same dataset as in Fig. 4[1] (a) and a validation cohort[2,3] (b), (c).

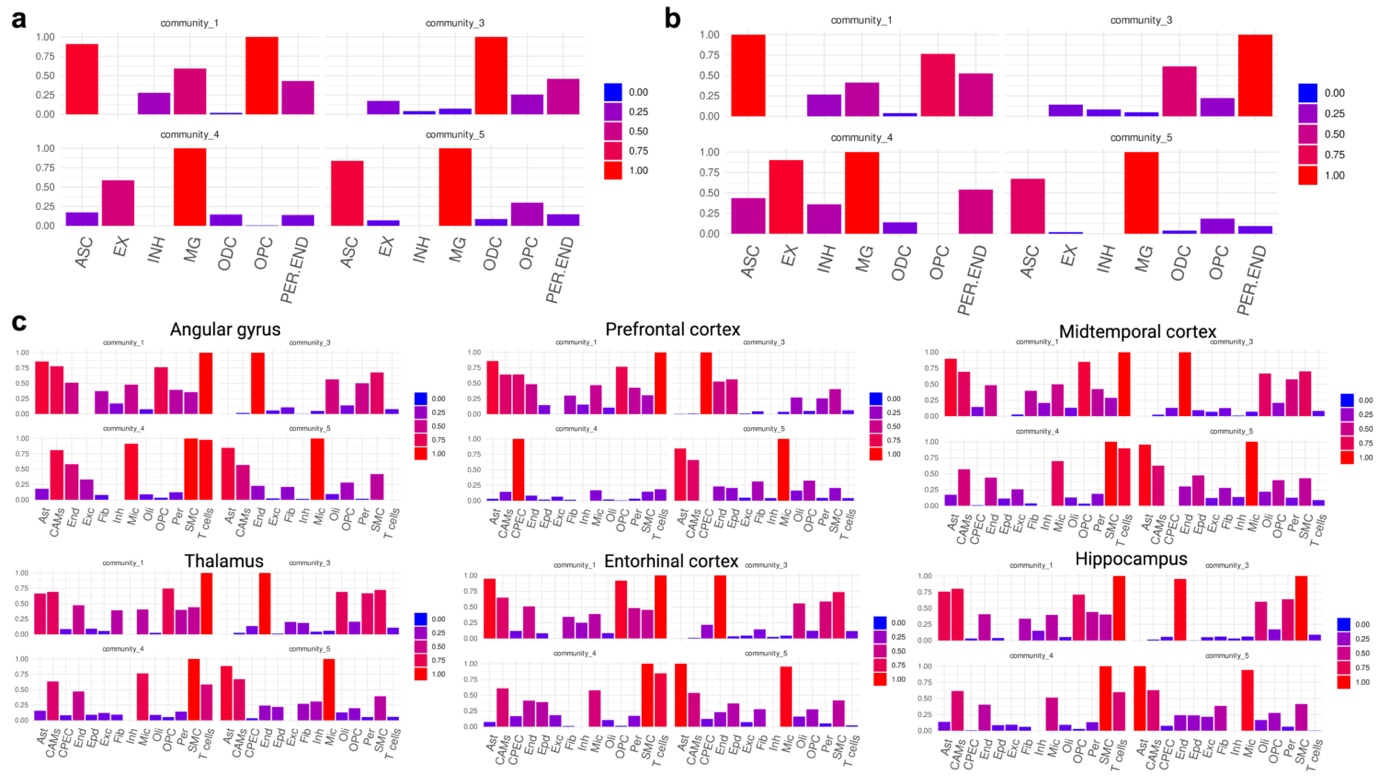

### **Supplementary Fig. S5 Characteristics of the ACT study Participants, related to Fig. 5**

The distribution of age (a), sex (b), CERAD score (c), and Braak score (d) of the ACT study participants differentiated between the no-dementia and dementia groups.

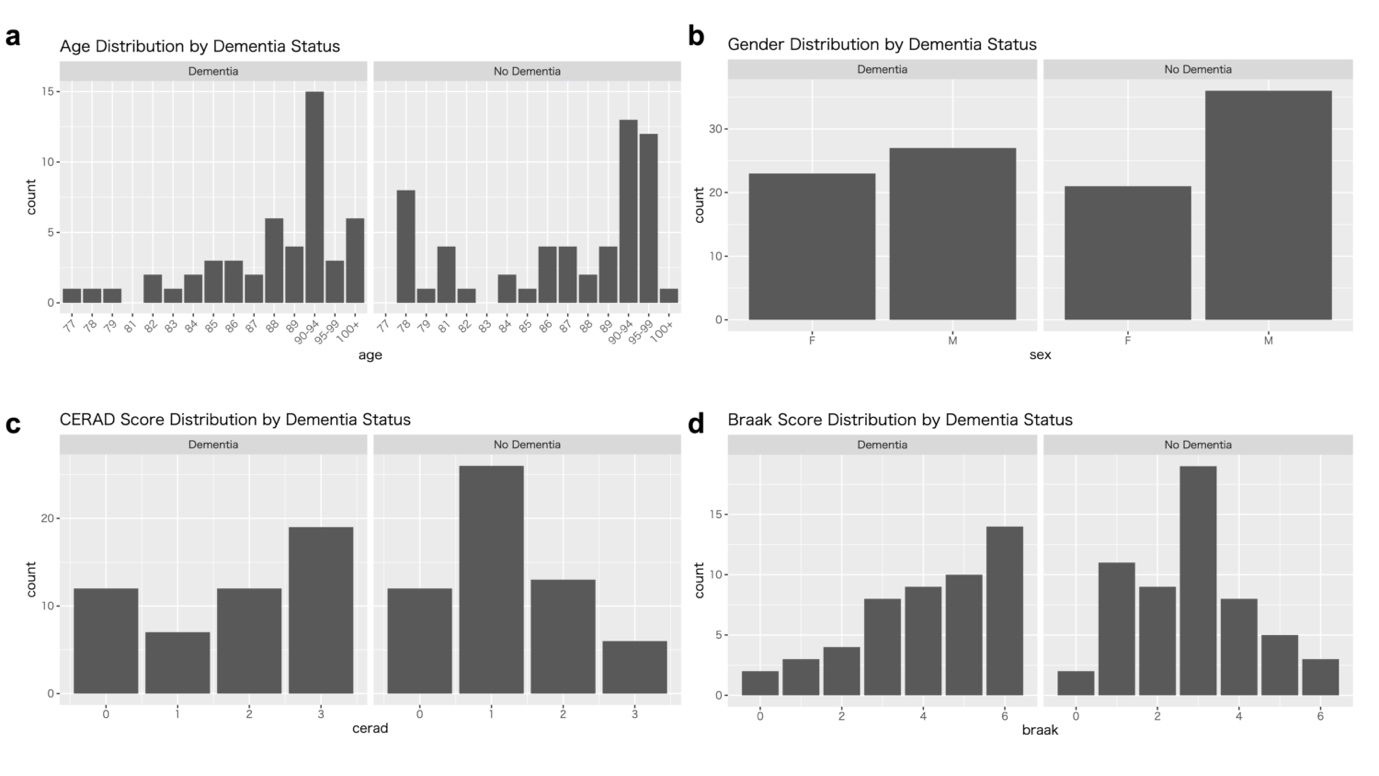

### **Supplementary Fig. S6 Cell Type Analysis for Control, Aβ+, and Aβ+Tau+ groups, related to Fig. 6**

Cell type compositions were compared among Control, Aβ+, and Aβ+Tau+ groups. Box plots displaying the distribution of cell type frequencies for each patient group. Results were analyzed using ANOVA, followed by Tukey's post-hoc test for pairwise comparisons when significant ANOVA results were observed.

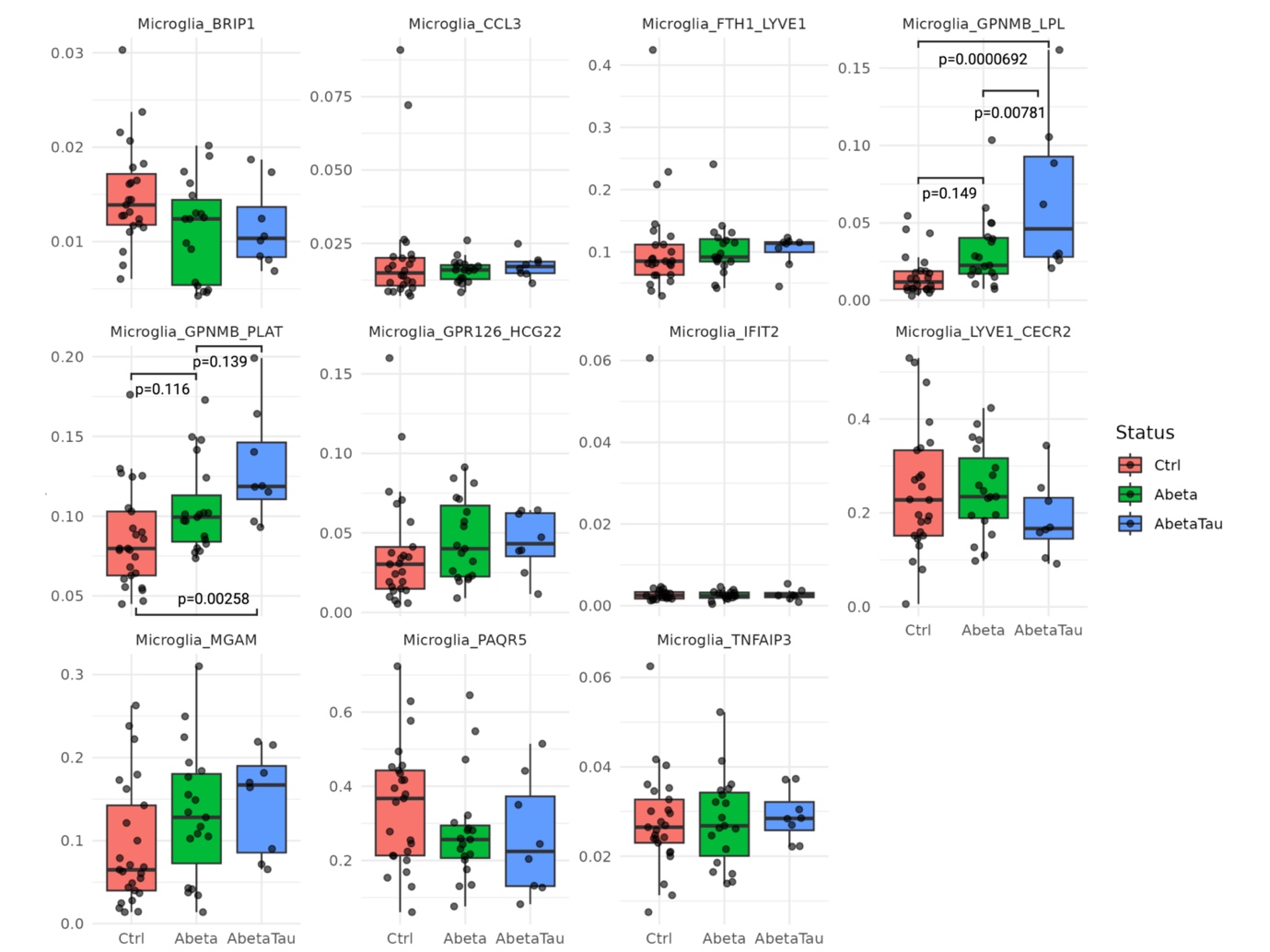

### **Supplementary Fig. S7 Trajectory analysis and pseudotime gene expression of candidate Aβ-tau-moderating factors, related to Fig. 6**

(a), (d) Slingshot trajectory inference applied to microglia revealed three lineages.

(b), (e) Pseudotime gene expression analysis of modulator candidate genes predicted using MINDy and GPNMB in a lineage. Gene expression levels are shown for the lineages that are highlighted in bold in (a) and (d), respectively.

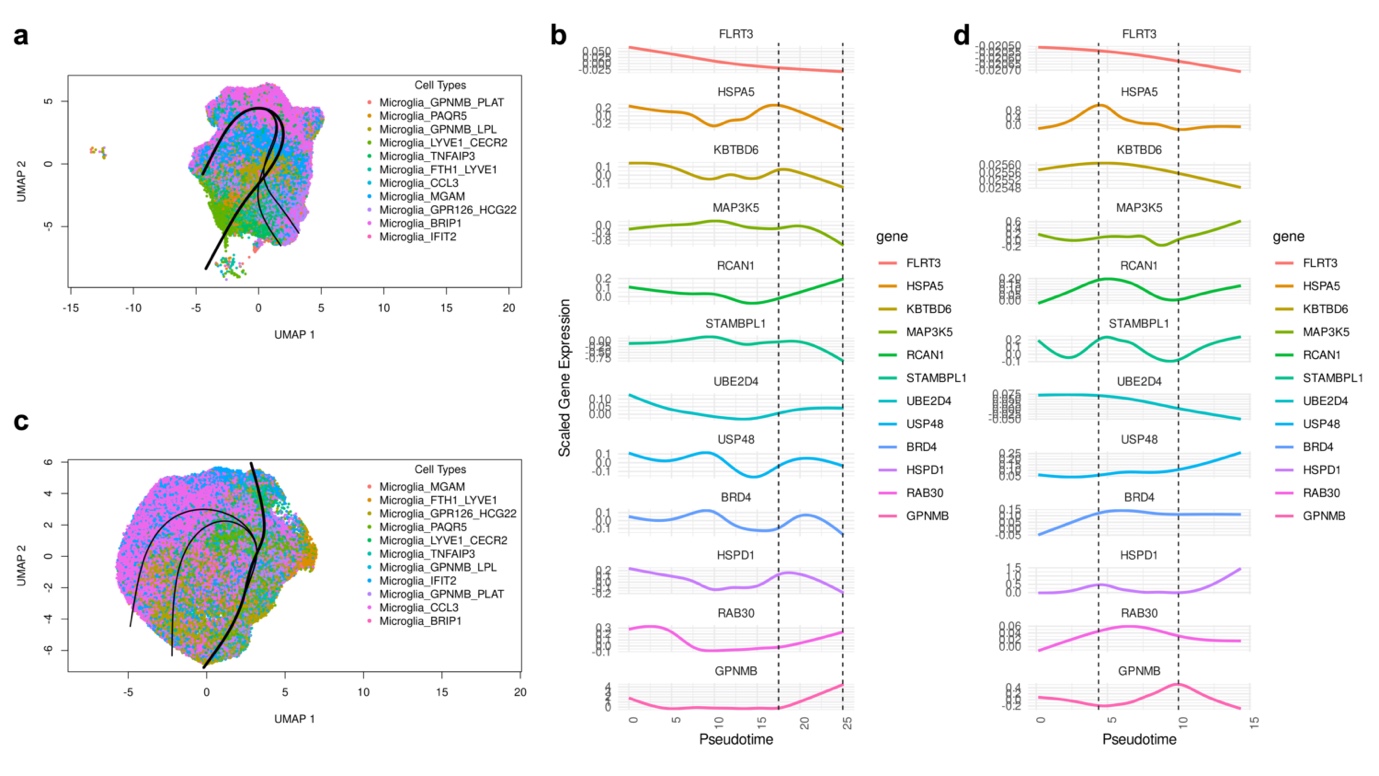

**Supplementary Table S1. Enrichment analysis for each module detected by clustering analysis, related to Fig. 2**

**Supplementary Table S2. Enrichment analysis for all subnetwork genes, related to Fig. 4**

**Supplementary Table S3. Enrichment analysis for each community within the subnetwork, related to Fig. 4**

**Supplementary Table S4. Linear interaction analyses using histopathological data with Tau as the dependent variable, and Aβ, GFAP, and IBA1 as the independent variables, related to Fig. 5.** For Tau, Tau2 antibody was used.

**Supplementary Table S5. Linear interaction analyses using histopathological data with Tau as the dependent variable, and Aβ, GFAP, and IBA1 as the independent variables, related to Fig. 5.** For Tau, AT8 antibody was used.

**Supplementary Table S6. Cell composition ratio of microglia subtypes in each pathological condition, related to Fig. S6**

**Supplementary Table S7. Linear interaction analyses using histopathological data with Tau as the dependent variable, and Aβ and GPNMB gene expression as the independent variables, related to Fig. 7.** For Tau, Tau2 antibody was used.

**Supplementary Table S8. Linear interaction analyses using histopathological data with Tau as the dependent variable, and Aβ and GPNMB gene expression as the independent variables, related to Fig. 7.** For Tau, AT8 antibody was used.

### **Supplementary Refences**

1. Morabito S, Miyoshi E, Michael N, Shahin S, Martini AC, Head E, et al. Single-nucleus chromatin accessibility and transcriptomic characterization of Alzheimer’s disease. Nat Genet [Internet]. 2021;53:1143–55. Available from: http://dx.doi.org/10.1038/s41588-021-00894-z

2. Sun N, Victor MB, Park YP, Xiong X, Scannail AN, Leary N, et al. Human microglial state dynamics in Alzheimer’s disease progression. Cell [Internet]. 2023;186:4386-4403.e29. Available from: http://dx.doi.org/10.1016/j.cell.2023.08.037

3. Gazestani V, Kamath T, Nadaf NM, Dougalis A, Burris SJ, Rooney B, et al. Early Alzheimer’s disease pathology in human cortex involves transient cell states. Cell [Internet]. 2023;186:4438-4453.e23. Available from: http://dx.doi.org/10.1016/j.cell.2023.08.005
